# Dynamic visualization of physiological CaMKII activity using sensitive FRET biosensors

**DOI:** 10.64898/2026.05.20.726522

**Authors:** Sohum Mehta, Nidhi A. Thaker, Kengo Adachi, Christopher Y. Ko, Bian Liu, Sai S. Divkaruni, Oguz C. Koc, Anne C. Lyons, Samantha A. Sanchez, Pauline Löffler, Yoshihisa Nakahata, Jody L. Martin, Jared L. Johnson, Tomer M. Yaron-Barir, Lewis C. Cantley, Martin J. Lohse, Andreas Bock, Donald M. Bers, Ryohei Yasuda, Rafael Fissore, Richard L. Huganir, Margaret M. Stratton, Jin Zhang

**Affiliations:** Department of Pharmacology, University of California San Diego, La Jolla, CA 92093, USA; Department of Biochemistry and Molecular Biology, University of Massachusetts, Amherst, MA 01003, USA; Max Planck Florida Institute for Neuroscience, Jupiter, FL 33458, USA; Department of Pharmacology, University of California Davis, Davis, CA 95616, USA; The Solomon H. Snyder Department of Neuroscience, The Johns Hopkins University School of Medicine, Baltimore, MD 21205, USA; Kavli Neuroscience Discovery Institute, The Johns Hopkins University School of Medicine, Baltimore, MD 21205, USA; Department of Neurology, The Johns Hopkins University School of Medicine, Baltimore, MD 21205, USA; Shu Chien-Gene Lay Department of Bioengineering, University of California San Diego, La Jolla, CA 92093, USA; Rudolf Boehm Institute of Pharmacology and Toxicology, Medical Faculty, University of Leipzig, 04107 Leipzig, Germany; Dana-Farber Cancer Institute, Harvard Medical School, Boston, MA 02115, USA; Department of Cell Biology, Harvard Medical School, Boston, MA 02115, USA; Department of Pediatrics, Boston Children’s Hospital, Harvard Medical School, Boston, MA 02115, USA; ISAR Bioscience Institute, 82152 Planegg, Munich, Germany; Institute of Pharmacology, University Medical Center of the Johannes Gutenberg-University Mainz, 55131 Mainz, Germany; Research Center for Immunotherapy (FZI), University Medical Center of the Johannes Gutenberg-University Mainz, 55131 Mainz, Germany; Department of Veterinary and Animal Sciences, University of Massacheusetts, Amherst, MA 01003, USA; Department of Chemistry and Biochemistry, University of California San Diego, La Jolla, CA 92093, USA; Moores Cancer Center, University of California San Diego, La Jolla, CA, 92093, USA

## Abstract

Calcium-calmodulin (CaM)-dependent protein kinase II (CaMKII) is a key mediator of complex physiological processes throughout the body, from the brain to the reproductive system, where CaMKII translates spatiotemporally dynamic calcium elevations into specific biological functions. Directly visualizing CaMKII activity dynamics in living cells using genetically encoded fluorescent biosensors can thus provide crucial insights into the molecular regulation of health and disease. Yet the ability to sensitively and specifically monitor endogenous CaMKII activity in physiologically relevant contexts is limited by the lack of sensors that can achieve robust, quantitative visualization of CaMKII responses. Here, we leveraged a recent serine/threonine kinome-wide substrate atlas to rationally engineer a powerful suite of Förster resonance energy transfer (FRET)-based CaMKII kinase activity reporters with high specificity, sensitivity, and signal-to-noise ratio. Using these biosensors, we were able to sensitively and robustly visualize endogenous CaMKII activity dynamics in both cultured cell lines and primary cells, including cardiomyocytes, oocytes, and neurons. We further utilized 2pFLIM imaging of organotypic hippocampal slices to quantitatively track LTP-induced CaMKII activity within single dendritic spines, highlighting a major advance in the study of physiological CaMKII signaling.

## INTRODUCTION

Ca^2+^/CaM-dependent protein kinase II (CaMKII) is a conserved serine/threonine kinase with major roles in transducing intracellular Ca^2+^ signals. Mammals express four major CaMKII variants^1,2^. The α and β isoforms are primarily found in the brain, particularly in postsynaptic densities within dendritic spines on excitatory neurons; δ is the principal cardiac isoform and is recruited to various subcellular structures within cardiomyocytes; and CaMKIIγ is ubiquitously expressed. Following activation via Ca^2+^/CaM binding and displacement of an autoinhibitory sequence, CaMKII can undergo autophosphorylation to enable CaM-independent activity^1,2^. The unique oligomeric structure of the CaMKII holoenzyme enables *in trans* autophosphorylation of individual subunits, implicating CaMKII as a form of molecular memory^3,4^ in numerous physiological processes, from activity-dependent synaptic plasticity^3^ to the cardiac force-frequency response^5,6^ and oocyte activation^7^, through frequency decoding of oscillatory Ca^2+^ signals. Illuminating the spatiotemporal dynamics of CaMKII activity is therefore crucial to unravelling these diverse and intricately coordinated functions and gaining insights into pathological mechanisms^3,8^.

Genetically encoded fluorescent biosensors are among the most powerful tools available for elucidating the spatiotemporal regulation of kinase signaling in cells^9^. Most CaMKII biosensors adopt a design originated by Camuiα^10^, in which full-length CaMKIIα is sandwiched between a pair of fluorescent proteins (FPs) such that Ca^2+^/CaM-induced conformational changes modulate FRET efficiency. Camui-like sensors are widely used to monitor CaMKII activation dynamics, including for specific variants^10–12^. However, their use necessitates overexpressing a single, full-length CaMKII variant and provides no information on the endogenous kinase, which comprises many variants. Instead, our recent FRET-based sensor of CaMKII activity (FRESCA)^13^ is designed to function as a surrogate substrate for CaMKII-mediated phosphorylation, enabling direct visualization of endogenous CaMKII activity in living cells. Given FRESCA’s small dynamic range, however, we set out to develop a CaMKII sensor capable of broader applications across a range of biological contexts to reveal greater insights into CaMKII signaling biology. Here, we report the development of rationally engineered, FRET-based CaMKII sensors with high selectivity and sensitivity, as well as expanded capability over existing tools. These biosensors enable robust and sensitive visualization of endogenous CaMKII activity dynamics across different biological systems, including cultured primary cardiomyocytes, oocytes, and neurons, as well as organotypic hippocampal slices, using fluorescence intensity- and lifetime-based approaches.

## RESULTS

### Development and characterization of FRESCA2

FRESCA follows the design of the previously described PKC activity reporter CKAR^14^, incorporating a forkhead-associated 2 (FHA2) domain and modified version of the *in vitro* CaMKII substrate “syntide-2”^15^ (syntide-Thr) as a CaMKII-sensitive molecular switch sandwiched between an mTurquoise2/Venus FRET pair^13^ (Fig. 1a). A phosphorylation-induced conformational change leads to decreased energy transfer, measured as a small increase in the cyan-over-yellow (C/Y) emission ratio (maximum ratio change [ΔR/R] = 2.16 ± 0.16%; mean ± s.e.m., n = 30 cells) (Fig. 1a), which was modestly enhanced (ΔR/R = 4.43 ± 1.8%, n = 31 cells) upon phosphatase inhibition with calyculin A (CalA, 25 nM; Supplementary Fig. 1a). We previously improved CKAR performance by replacing the FHA2 domain with FHA1^16^ and thus applied a similar strategy to FRESCA, swapping syntide-Thr into a sensor backbone containing FHA1 and a Cerulean/cpVenus[E172] FRET pair^17^ (Fig. 1b). We also incorporated a Gly-to-Asp substitution at +3 relative to the phospho-acceptor Thr in the substrate (S1; PLARALTVA<u>D</u>LPGKK) to match FHA1 binding preferences^18^ (Fig. 1b).

**Figure 1.**
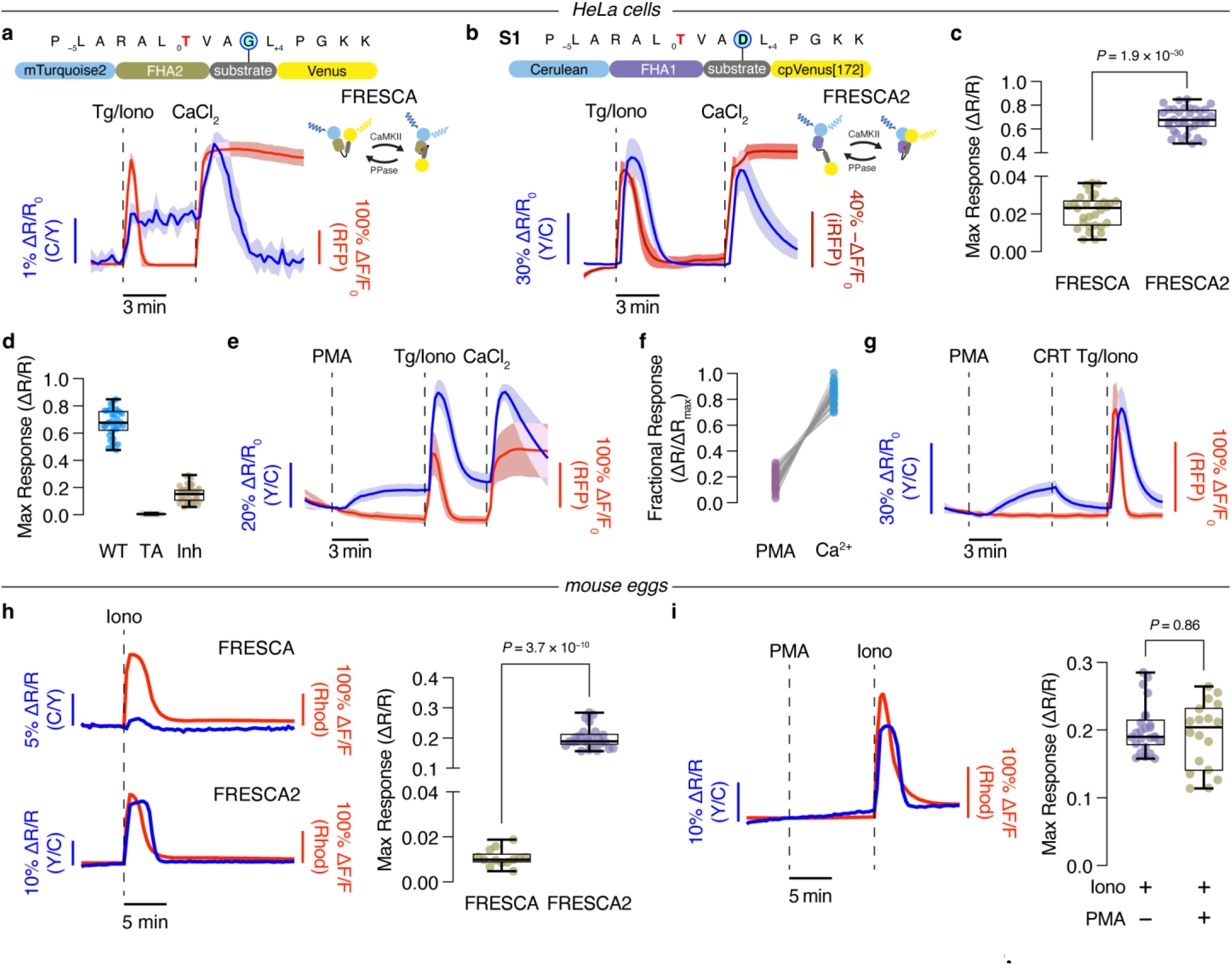
Improving FRESCA performance through domain substitution. **a-b**, Representative average timecourses of (**a**) FRESCA C/Y (blue, n=8 cells) or (**b**) FRESCA2 Y/C (blue, n=13 cells) ratio and RCaMP1d (**a**, red) or NIR-GECO2G (**b**, dark red) intensity change in HeLa cells stimulated with 1 μM thapsigargin plus 2 μM ionomycin (Tg/Iono) followed by 5 mM CaCl_2_. Biosensor schematics shown above. **c**, Maximum Ca^2+^-stimulated FRESCA (n=30 cells) or FRESCA2 (n=26 cells) ratio change (ΔR/R). **d**, Maximum Ca^2+^-stimulated ratio changes (ΔR/R) from FRESCA2 (WT, n=36 cells) or FRESCA2 T/A (TA, n=40 cells) plus mCherry, or FRESCA2 plus CaMKip-mCherry (Inh, n=40 cells). **e**, Representative averaged timecourse of FRESCA2 ratio (blue) and RCaMP1d intensity (red) change in response to 50 ng mL^−1^ phorbol 12-myristate 13-acetate (PMA), Tg/Iono, and CaCl_2_. n = 13 cells. **f**, Fractional PMA- and Ca^2+^-stimulated FRESCA2 responses. n = 42 cells. **g**, Representative averaged timecourses of FRESCA2 ratio (blue) and RCaMP1d intensity (red) change in cells treated with 5 μM CRT0066101 (CRT) after PMA, followed by Tg/Iono and CaCl_2_. n = 9 cells. Solid lines (**a**, **b**, **e**, and **f**) indicate means; shaded areas, s.d. Box-and-whisker plots (**c**, **d**, **h**, and **i**) show median, quartiles, min, and max. **h**, Left: Representative single-cell timecourses of FRESCA C/Y (top, n=13 eggs) or FRESCA2 Y/C (bottom, n=16 eggs) ratio (blue curves) and Rhod-2 intensity (red curves) in oocytes stimulated with 500 nM Iono. Right, maximum Iono-induced ratio changes (ΔR/R) from FRESCA (n=13 eggs) or FRESCA2 (n=16 eggs). **i**, Left, representative single-cell timecourse of FRESCA2 ratio (blue curve, n=18 eggs) and Rhod-2 intensity (red curve) in oocytes stimulated with 1 µM PMA followed by 500 nM Iono. Right, maximum Iono-induced FRESCA2 response with (+, n=18 eggs) or without (−, n=25 eggs) PMA pre-treatment. Timecourses represent (**a**, **b**, **e**, **h**, **i**) 3 or (**g**) 4 independent experiments each. Data in **c**, **d**, **f**, **h**, and **i** pooled from 3 independent experiments each. Unpaired, two-tailed Student’s t-test (**c**) or Mann-Whitney U-test (**h**, **i**). See also Supplementary Figs. 1-3.

HeLa cells expressing this FRESCA2 construct exhibited a robust increase in the yellow-over-cyan (Y/C) emission ratio, which closely mirrored the response from a co-expressed Ca^2+^ indicator^19^, upon addition of 1 μM thapsigargin plus 2 μM ionomycin (Tg/Iono) to promote endoplasmic reticulum (ER) store release (Fig. 1b). Subsequent addition of 5 mM CaCl_2_ elicited an additional response (Fig. 1b), with a maximum ΔR/R of 67.6 ± 1.7% (n = 36 cells), representing a >30-fold performance enhancement over FRESCA (*P* = 1.93 × 10^−30^; Fig. 1c and Supplementary Fig. 1b). Addition of CalA prior to either Tg/Iono or CaCl_2_ treatment led to sustained elevation of the FRESCA2 emission ratio (Supplementary Fig. 1c-e), while a Thr-to-Ala phospho-acceptor mutant sensor (FRESCA2 T/A) did not respond to Ca^2+^ (Supplementary Fig. 2a), confirming that the observed FRET changes correspond to phosphorylation and dephosphorylation of the biosensor.

However, when we blocked CaMKII activity by overexpressing an mCherry-tagged CaMKII inhibitor peptide (CaMKip) based on autocamtide inhibitory peptide 2^20^, we observed a residual Tg/Iono-stimulated response from FRESCA2 (Fig. 1d and Supplementary Fig. 2b). Crucially, another member of the CAMK family, Protein Kinase D (PKD), can also phosphorylate syntide-2 *in vitro*^21^. Using the FRET-based PKD sensor DKAR^22^ in HeLa cells, we confirmed that PKD activity is induced by Tg/Iono stimulation, in addition to phorbol 12-myristate 13-acetate (PMA) (Supplementary Fig. 2c). We further observed a clear response from FRESCA2 upon PMA stimulation (Fig. 1e), which comprised a modest fraction of the sensor’s total dynamic range (Fig. 1f) and was reversed by the PKD inhibitor CRT0066101 (CRT)^23^ without blocking Ca^2+^-induced responses (Fig. 1g). PKD inhibition also suppressed the residual Tg/Iono-induced FRESCA2 response in HeLa cells co-expressing CaMKip (ΔR/R = 15.3 ± 0.9% vs. 8.15 ± 0.67%, n = 39 cells each; *P* = 2.16 × 10^−7^) (Supplementary Fig. 2b) and slightly reduced the maximum Tg/Iono-stimulated FRESCA2 response (ΔR/R = 67.6 ± 1.7%, n = 36 cells vs. 57.7 ± 1.7%, n = 28 cells; *P* = 1.38 × 10^−4^) (Supplementary Fig. 2d).

Notably, we did not previously observe cross-phosphorylation of FRESCA in mouse eggs^13^, potentially due to either 1) increased sensitivity of FRESCA2 revealing weak phosphorylation by PKD or 2) cell-type-specific differences in PKD activity. Similar to HeLa cells, mouse eggs expressing FRESCA2 exhibited an 18-fold greater response to Iono stimulation than cells expressing FRESCA (ΔR/R = 20.2 ± 4.6%, n = 25 eggs vs. 1.1 ± 0.4, n = 13 eggs; *P* = 3.7 × 10^−10^) (Fig. 1h). Yet stimulating these cells with PMA yielded no FRESCA2 response (Fig. 1i and Supplementary Fig. 3), in contrast to our results in HeLa cells (Fig. 1e). Thus, despite the potential for nonspecific PKD phosphorylation in some cell types, FRESCA2 offers greatly enhanced performance over the original FRESCA and should facilitate more sensitive visualization of CaMKII activity dynamics, for example, in mouse eggs.

### Rationally engineering a CaMKII sensor with increased specificity

Individual kinases maintain specificity by recognizing characteristic sequence motifs surrounding their target residues, which corresponds to the substrate sequence in a kinase biosensor. Given the potential for cross-phosphorylation by PKD, we therefore sought to re-engineer the FRESCA2 substrate sequence to increase selectivity. Previously, rational engineering of a *de novo* substrate for a single kinase^24^ was enabled using peptide libraries^25^ in which a 10-residue region spanning 5 positions N-terminal (P_-1_ to P_-5_) and 4 positions C-terminal (P_+1_ to P_+4_) to the phosphosite (P_0_) is systematically scanned with different amino acids. However, engineering a selective substrate that discriminates between related kinases requires a new strategy. Based on a recent kinome-wide atlas^26^ in which the substrate preferences of >300 Ser/Thr kinases were evaluated using scanning peptide libraries and quantitatively summarized as positional site scoring matrices (PSSMs), we hypothesized that comparing CaMKII and PKD substrate preferences would yield an optimized substrate sequence that eliminates PKD phosphorylation.

To quantitatively compare the amino acid preferences of CaMKIIα and PKD1, we devised an analytical strategy that should be broadly generalizable to multiple kinases and instances of substrate cross-phosphorylation. After first extracting PSSM data for our kinase of interest (KOI; CAMK2A) and kinase of detriment (KOD; PRKD1) from the kinome atlas dataset, we generated a “selectivity matrix” (see Methods) by dividing the KOI and KOD matrices (i.e., CAMK2A/PRKD1) (Fig. 2a). We then normalized the KOI matrix against the selectivity matrix to generate a final PSSM that balances sensitivity for the KOI and selectivity over the KOD. The resulting substrate matrix allowed us to identify specific residue substitutions in the FRESCA2 substrate (S1) that were predicted to influence favorability towards CaMKII versus PKD (Fig. 2a). Predicted substrate modifications were examined using an online tool (https://kinase-library.phosphosite.org/kinase-library/)^26^ to determine their Log2 Score Rank, denoting how well a particular sequence (P_−5_ to P_+4_) will be phosphorylated by each of the kinases in the kinome atlas dataset. Consistent with our HeLa cell data, numerous kinases, including PRKD1, rank higher than CAMK2A in phosphorylating S1 (Supplementary Table 1).

**Figure 2.**
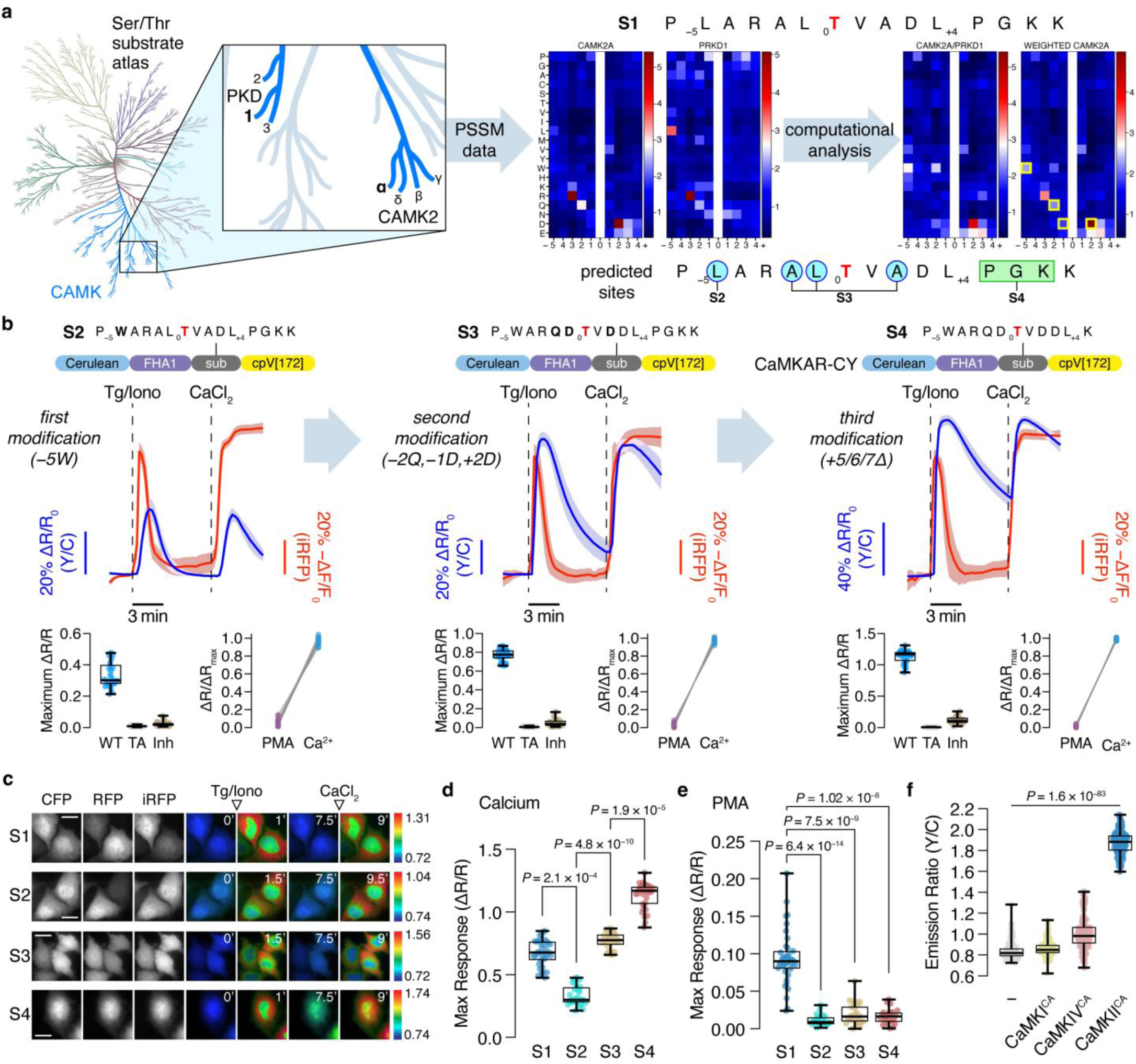
Rational design of CaMKAR-CY with enhanced selectivity. **a**, CaMKIIα (CAMK2A) and PKD1 (PRKD1) data extracted from a kinome-wide atlas were mathematically transformed to predict discriminatory substrate residues (yellow boxes). Substrate modifications are indicated below. **b**, Top, domain schematics and representative averaged timecourses showing Y/C ratio and intensity changes from HeLa cells expressing S2 (left, n = 6 cells), S3 (middle, n = 12 cells), or S4 (right, n = 8 cells; CaMKAR-CY) plus NIR-GECO2G and stimulated with Tg/Iono and CaCl_2_. Representative of 3 independent experiments each. Bottom left, maximum Ca^2+^-stimulated ratio changes (ΔR/R) from HeLa cells expressing S2 (left), S3 (right), or S4 (right) (WT) or the T/A mutant (TA) co-transfected with mCherry or CaMKip-mCherry (Inh). n = 27, 41, and 33 (S2); 33, 36, and 28 (S3); and 35, 39, and 29 (S4) cells. Bottom right, fractional responses upon PMA and Ca^2+^ stimulation. n = 24, 26, and 31 cells. **c**, Y/C ratio (pseudocolor) images of S1 (FRESCA2), S2, S3, and S4 (CaMKAR-CY) upon Tg/Iono and CaCl_2_ stimulation. CFP, RFP, and iRFP images show construct expression. Arrowheads indicate stimulation. Representative of 3 independent experiments. Scale bars, 10 μm. **d**-**e**, Maximum Ca^2+^ (**d**)- or PMA (**e**)-stimulated ratio changes from S1, S2, S3, and S4. Data from 36, 27, 33, and 35 (**d**) and 42, 24, 26, and 22 (**e**) cells. **f**, Raw Y/C ratios from HeLa cells expressing CaMKAR-CY plus mCherry or mCherry-tagged constitutively active CaMKI, CaMKIV, or CaMKII. Data from 148, 127, 137, and 145 cells. Kruskal-Wallis test followed by Dunn’s multiple comparisons test. Solid lines in **b** indicate the mean; shaded areas, s.d. Box-and-whisker plots in **b** and **d**-**f** show median, quartiles, min, and max. Data from (**b**, **d**, **e**) 3 or (**f**) 4 independent experiments. See also Supplementary Figs. 4-6.

Our analysis first highlighted the P_−5_ site, where incorporating Trp (−5W) was predicted to strongly disfavor PKD phosphorylation. The resulting sequence (S2; P_−5_<u>W</u>ARALTVADL_+4_PGKK) (Fig. 2b) yielded a substantially decreased Log2 Score Rank for PKD phosphorylation versus S1 (Supplementary Table 1). HeLa cells expressing an S2-containing biosensor exhibited clear, Ca^2+^-stimulated Y/C emission ratio changes (maximum ΔR/R = 33.5 ± 1.5%, n = 27 cells) (Fig. 2b-e), whereas we observed no responses in cells expressing a T/A-mutant S2 biosensor (ΔR/R = 0.84 ± 0.06%, n = 41 cells) or co-expressing mCherry-CaMKip (ΔR/R = 2.45 ± 0.29%, n = 33 cells) (Fig. 2b and Supplementary Fig. 4). Direct stimulation with PMA also elicited no response (ΔR/R = 1.14 ± 0.15%, n = 24 cells; *P* = 6.42 × 10^−14^ vs. S1) (Fig. 2b,e and Supplementary Fig. 4).

Despite having a similar Log2 Score Rank as S1 for CaMKII (Supplementary Table 1), the Ca^2+^-stimulated response of the S2 biosensor was significantly lower (*P* = 6.73 × 10^−5^; Fig. 2d). An S2 peptide also showed poor phosphorylation *in vitro* (Supplementary Fig. 5). We subsequently predicted that converting P_−2_, P_−1_, and P_+2_ to Gln, Asp, and Asp (−2Q/−1D/+1D) would greatly improve CaMKII-mediated phosphorylation. Indeed, CAMK2A was the top-ranked kinase for this new sequence (S3; P_−5_WAR<u>QD</u>TV<u>D</u>DL_+4_PGKK), while maintaining a low Log2 Score Rank for PRKD1 (Supplementary Table 1), which translated into improved phosphorylation *in vitro* (Supplementary Fig. 5). Consistent with these results, S3 biosensor-expressing HeLa cells showed much higher Ca^2+^-stimulated emission ratio changes (ΔR/R = 77.3 ± 1.1%, n = 33 cells) than cells expressing the S2 biosensor (*P* = 4.78 × 10^−10^; Fig. 2b-d). The S3 biosensor also retained good selectivity over PKD, showing little response to Ca^2+^ stimulation when co-expressed with mCherry-CaMKip (ΔR/R = 5.24 ± 0.803%, n = 28 cells) or when directly stimulated with PMA (ΔR/R = 1.98 ± 0.28%, n = 26 cells; *P* = 7.52 × 10^−9^ vs S1) (Fig. 2b,e and Supplementary Fig. 4).

Finally, we truncated sites P_+5_ to P_+7_ to obtain a “minimal” substrate, similar to other kinase sensors^17,24,27^, without interfering with the core sequence defined by PSSM data or altering junctions with other sensor domains. This substrate (S4; P_−5_WARQDTVDDL_+4_K) could not be scored, as the modification falls outside the PSSM data, and behaved similarly to S3 *in vitro* (Supplementary Fig. 5). Nevertheless, the S4 biosensor showed even further enhancement, with an average ΔR/R of 113 ± 2%, (n = 35 cells) upon Ca^2+^ stimulation in HeLa cells (Fig. 2b-d) and no apparent phosphorylation by PKD (Fig. 2e, Supplementary Fig. 4). The S4 biosensor also showed minimal response to constitutively active CaMKI or CaMKIV (Fig. 2f), along with similar response amplitude and faster onset kinetics versus a recently reported single-FP based CaMKII activity reporter^27^ (Supplementary Fig. 6). Thus, we were successfully able to apply recently published substrate consensus sequence information of the human kinome to rationally engineer a sensitive and specific C/Y-FRET-based CaMKII kinase activity reporter (CaMKAR-CY).

### Visualizing endogenous CaMKII activity in cardiac myocytes

CaMKII is a central and critical regulator of many key processes and functions in cardiac myocytes, such as excitation-contraction coupling, Ca^2+^ handling, ionic currents, and transcription^28,29^. Persistent overactivation of CaMKII, exacerbated by numerous post-translational modifications^30^, can lead to cardiac disease, such as heart failure and arrhythmias. The ability to monitor CaMKII signaling in cardiac myocytes is therefore critically important but has previously been limited to indirect conformational sensors or destructive methods to probe target phosphorylation. CaMKAR-CY overcomes these limitations by enabling the direct and dynamic measurement of CaMKII target phosphorylation in real time, reporting endogenous CaMKII activity in its native intracellular cardiac myocyte environment (Fig. 3). When expressed in isolated rabbit ventricular myocytes by adenoviral infection, CaMKAR-CY is present throughout the myocyte (Fig. 3a), and a robust functional FRET response was confirmed via acceptor photobleach-induced donor enhancement (Supplementary Fig. 7a). When resting myocytes were electrically paced at 1 Hz to evoke action potentials and concomitant Ca^2+^ transients that drive Ca^2+^/CaM dependent activation of CaMKII, the normalized FRET ratio (R/R_0_, where R = F_DA_/F_DD_) gradually increased over 10 min in control but rose much faster in myocytes preincubated and superfused with the β-adrenergic agonist isoproterenol (Iso), peaking in 2 min (Fig. 3b, left). Consistent with enhanced FRET, both an increase in acceptor YFP fluorescence (F_DA_) and a decrease (or quenching) of the donor CFP signal (F_DD_) were observed (Fig. 3b, right). The rise in CaMKAR-CY signal is what is expected for pacing-induced CaMKII activation in rabbit ventricular myocytes, as revealed previously using the CaMKII activation reporter, Camui, whose response is also accelerated by Iso^31^. The decline in Iso-treated CaMKAR-CY signal from 2 to 10 min is not atypical for the overall myocyte Ca^2+^-related functional response to Iso^32,33^ and may reflect slower activation of parallel pathways that limit acute inotropic effects, including activation of phosphatases, ion channels, and transporters.

**Figure 3.**
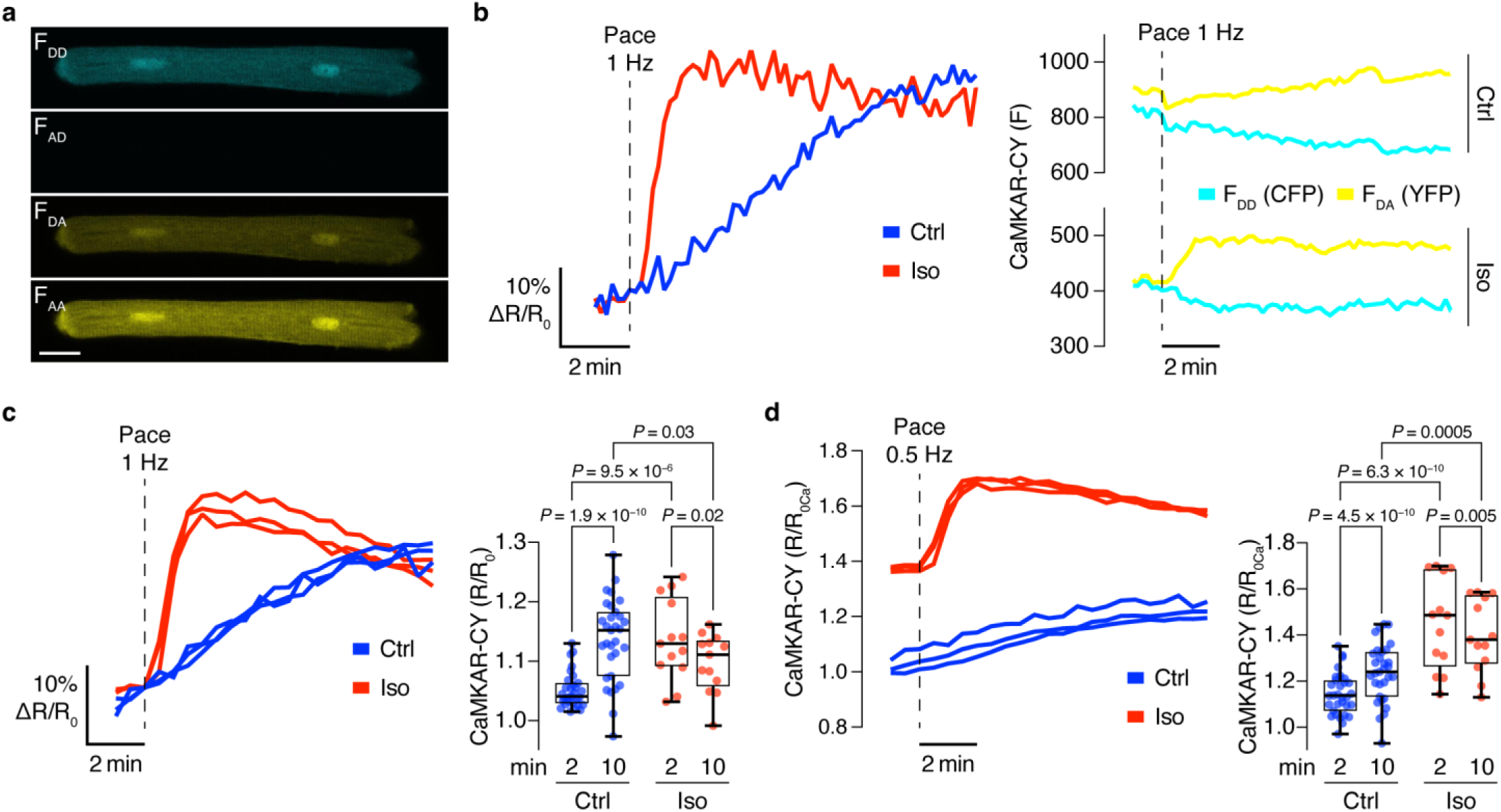
CaMKII activity imaging in cardiac myocytes. **a**, Representative fluorescence (F) image showing CaMKAR-CY expression in a rabbit ventricular myocyte. DD: Donor excitation, Donor emission; AD: Acceptor excitation, Donor emission; DA (FRET): Donor excitation, Acceptor emission; AA: Acceptor excitation, Acceptor emission. Scale bar, 20 μm. **b**, Representative timecourses of normalized CaMKAR-CY emission ratio (R/R_0_) upon 1 Hz field-stimulation pacing in control (Ctrl, blue)- or isoproterenol (Iso, 100 nM, incubation ∼3 min; red)-stimulated cardiac myocytes (left) and corresponding individual CFP (F_DD_) and YFP (F_DA_) intensity traces (right). **c**, Left, representative timecourses of individual normalized emission ratios (R/R_0_) at 0.5 Hz field-stimulation pacing onset under Ctrl and Iso treatment. Right, quantification of normalized emission ratios (R/R_0_) at 2 and 10 min after pacing onset under Ctrl and Iso treatment. **d**, Left, representative timecourses of individual emission ratios (R) depicted in **c** normalized to the median emission ratio of myocytes at 0 mM Ca^2+^ (R_0Ca_=0.548; Extended Data Fig. 6c). Right, quantification of normalized emission ratios (R/R_0Ca_) at 2 and 10 min after pacing onset under Ctrl and Iso treatment. Ctrl: n_rabbits_=3, n_myocytes_=31; Iso: n_rabbits_=2, n_myocytes_=13. Repeated measures two-way ANOVA mixed-effect model followed by Fisher’s Least Squares Difference with single-pooled variance for multiple comparisons post hoc analysis (**c, d**). Box-and-whisker plots in **c** and **d** show median, quartiles, min, and max. See also Supplementary Fig. 7.

While the CaMKAR-CY responses during pacing rose more quickly with Iso than in control conditions, we also expected a much higher peak with Iso. Indeed, the absolute R values (not normalized to R_0_) were higher in the presence of Iso vs. control (0.71 vs. 0.60, Supplementary Fig. 7d) indicating that CaMKII was already more active with Iso versus control in quiescent myocytes, in agreement with prior measurements with Camui^31^. Untreated resting cardiac myocytes have also previously been reported to exhibit significant basal CaMKII activity^34^, prompting us to test whether CaMKAR-CY is able to sense partial CaMKII target phosphorylation in the quiescent, resting state. We therefore preincubated ventricular myocytes in Ca^2+^-free solution for 30 min, to minimize CaMKII activation, before introducing the 1.8 mM [Ca^2+^]_o_ solution used as control in Fig. 3 (Supplementary Fig. 7e), which resulted in a gradual rise in the CaMKAR-CY signal by 20% over 25 min. Renormalizing the CaMKAR-CY response curve (Fig. 3c) to the Ca^2+^-free baseline signal (R_0Ca_) thus revealed a larger dynamic range (>60% increase in R/R_0Ca_), comparable to the values observed in HeLa cells (Fig. 2).

Thus, CaMKAR-CY can be effectively implemented as a genetically encoded molecular reporter of CaMKII target phosphorylation in live cardiac myocytes. The signal responses to both physiological and pharmacological experimental interventions are robust and allow for the direct measurement of the physiological range of dynamic CaMKII activity in real time. These qualities make CaMKAR-CY valuable for uncovering the mechanistic role of CaMKII in regulating essential processes in cardiac myocytes and can help reveal valuable insights into understanding CaMKII-mediated mechanisms of cardiac physiology and disease.

### Monitoring oscillatory CaMKII activity in mouse eggs

The fusion of a sperm and egg induces cytosolic Ca^2+^ elevations that serve as a universal trigger for key early events of egg activation and embryo development^35^. Downstream CaMKII activation is essential for transmitting fertilization-induced Ca^2+^ signals^36^, and female mice that lack CaMKII expression are sterile despite their eggs showing normal Ca^2+^ responses^37^. Tools to study CaMKII activity in living cells could thus provide important insights into the molecular regulation of fertilization. In mouse eggs transiently expressing CaMKAR-CY, we found that Iono stimulation yielded large, rapid Y/C emission ratio increases that closely corresponded to transient Ca²⁺ elevations observed using the fluorescent dye Rhod-2 (Fig. 4a, b, Supplementary Fig. 8a, b). Eggs expressing a T/A mutant sensor or co-expressing CaMKIIN1^38^ showed no responses (Supplementary Fig. 8c, d). CaMKAR-CY exhibited a 29.7 ± 1.5% (n = 18 eggs) emission ratio change in response to Iono stimulation compared with a 20.2 ± 0.7% (n = 25 eggs; *P* = 1.6 × 10^−6^) for FRESCA2 (Fig. 4c and Supplementary Fig. 8e, f), highlighting its improved performance in mouse eggs. Thus, CaMKAR-CY should enable highly sensitive and specific detection of physiological CaMKII activity changes.

**Figure 4.**
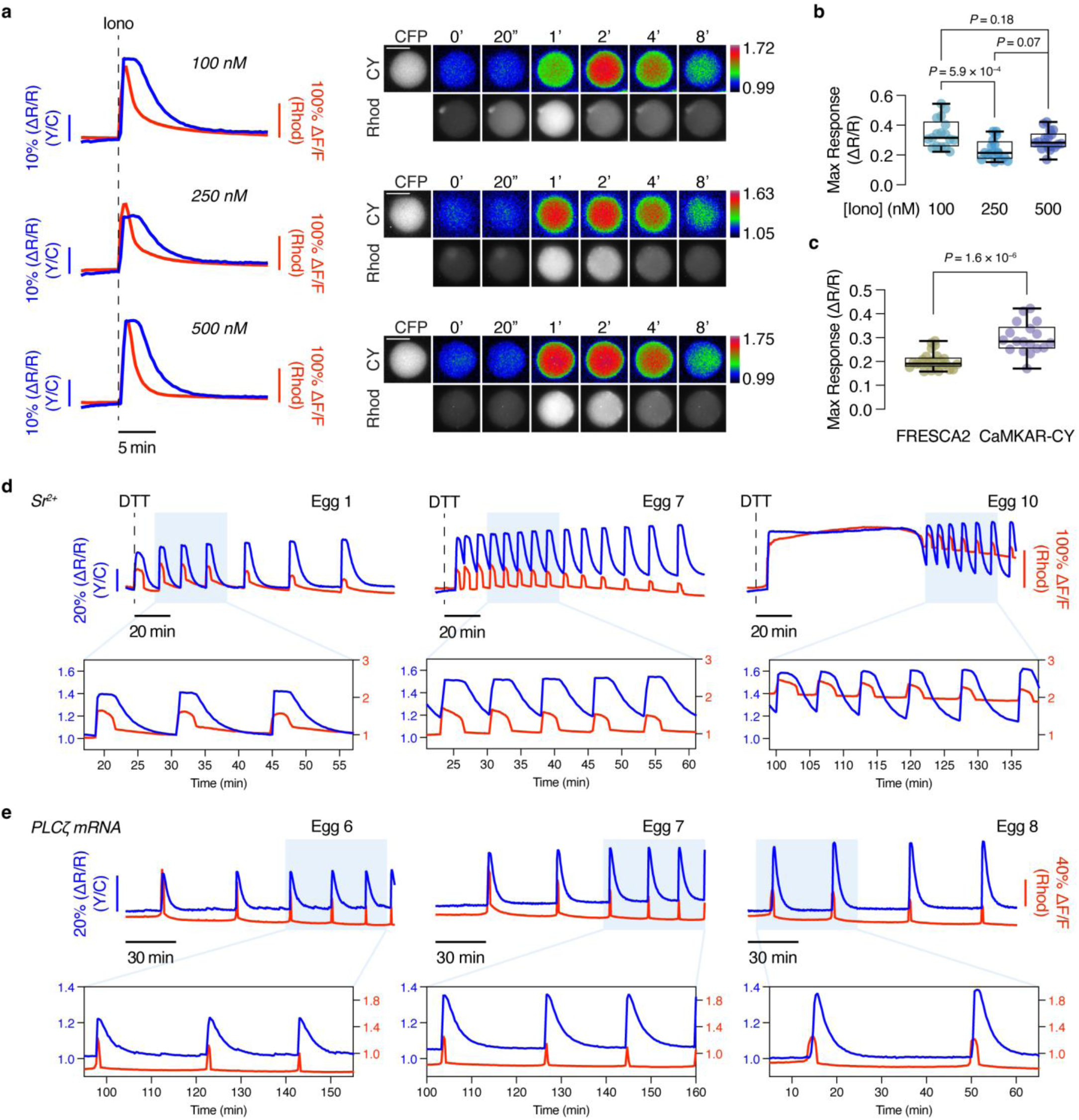
Visualizing oocyte CaMKII activity dynamics. **a**, Left, representative timecourses of CaMKAR-CY Y/C ratio (blue) and Rhod-2 fluorescence intensity (red) changes in oocytes stimulated with 100 nM (n=17 eggs), 250 nM (n=16 eggs), and 500 nM (n=18 eggs) Iono. Data are representative of 3 independent experiments each. Right, representative images showing CaMKAR-CY (CY) emission ratio (pseudocolor) and Rhod-2 (Rhod) fluorescence in oocytes at the indicated times before and after Iono addition. Warmer colors indicate higher ratios. CFP channels images show CaMKAR-CY expression. Scale bars, 50 μm. **b**, Maximum Iono-stimulated ratio change (ΔR/R) in CaMKAR-CY-expressing oocytes. n=17 (100 nM), 16 (250 nM), and 18 (500 nM) eggs from 3 independent experiments. **c**, Maximum Iono-stimulated ratio changes (ΔR/R) from FRESCA2 (n=25 eggs) and CaMKAR-CY (n=18 eggs) in oocytes stimulated with 500 nM Iono. Data from 3 independent experiments each. Box-and-whisker plots show the median, quartiles, min, and max. Ordinary one-way ANOVA(**b**) or Mann-Whitney U-test (**c**). **d**-**e**, Representative timecourses showing CaMKAR-CY emission ratio (blue) and Rhod-2 fluorescence intensity (red) changes from selected oocytes exhibiting oscillatory responses upon (**d**) incubation with 10 mM Sr²⁺ and addition of 1 mM DTT (n=20 eggs from 4 independent experiments) or (**e**) injection of PLCζ mRNA (n=19 eggs from 4 independent experiments). Insets display zoomed-in views of peaks within the shaded regions. See also Supplementary Figs. 8 and 9.

In mammals, fertilization is marked by the onset of prolonged oscillations in cytosolic Ca^2+^ concentrations. Previous studies established a direct relationship between CaMKII activity increases and the onset of Ca^2+^ oscillations^7,39–41^ but were limited to the first hour post-fertilization. Thus, a complete picture of CaMKII activity dynamics in response to fertilization-induced Ca^2+^ oscillations is lacking. Given the robust performance of CaMKAR-CY, we next sought to visualize endogenous CaMKII activity in response to oscillatory Ca²⁺ signaling. Application of extracellular Sr^2+^ can induce parthenogenetic activation of mouse eggs via TRPV3-mediated influx^42^, leading to robust oscillatory signaling. In CaMKAR-CY-expressing eggs incubated in media containing 10 mM Sr²⁺, treatment with 1 mM DTT triggered a large, variable initial rise in Rhod-2 fluorescence intensity, likely reflecting combined Sr^2+^/Ca^2+^ elevation, followed by robust and sustained oscillations (Fig. 4d). We also observed clear and striking emission ratio increases from CaMKAR-CY that were synchronized with Rhod-2 fluorescence peaks, indicating robust, oscillatory CaMKII activity (Fig. 4d).

Sr²⁺-induced oscillations occur at a higher frequency and exhibit broader peaks than native Ca²⁺ oscillations observed *in vivo* during fertilization, wherein PLCζ released from the fused sperm induces IP_3_-mediated release of ER Ca^2+^ stores^43^. Indeed, directly injecting PLCζ mRNA into CaMKAR-CY-expressing eggs to more closely simulate fertilization yielded slower Ca^2+^ oscillations, characterized by smaller, more rapid Ca^2+^ transients (Fig. 4e). Nevertheless, CaMKAR-CY effectively responded to these oscillations, reaching a similar amplitude as seen with Sr^2+^/DTT treatment (Fig. 4e). Peak-to-peak CaMKAR-CY responses were also highly consistent under both Sr^2+^/DTT- and PLCζ-stimulated conditions (Supplementary Fig. 8d, e) and steady throughout the Ca^2+^ oscillations. Importantly, the CaMKAR-CY emission ratio increased in response to the first Ca^2+^ peak during both Sr^2+^/DTT- and PLCζ-induced oscillations, in striking contrast to delayed responses seen previously with Camuiα and FRESCA^13^. These results underscore the sensitivity of CaMKAR-CY to visualize physiological CaMKII signaling and indicate that CaMKII activity is immediately triggered by and robustly maintained throughout fertilization-induced Ca^2+^ oscillations^13^. CaMKAR-CY is thus a powerful tool for monitoring CaMKII kinase activity dynamics and providing new insights into the regulation of Ca²⁺ signaling during fertilization.

### Sensitive detection of endogenous neuronal CaMKII activity

CaMKII plays important roles in synaptic plasticity, which is widely regarded as the cellular correlate of learning and memory^3^. In primary rat hippocampal neurons expressing CaMKAR-CY, Iono stimulation induced a robust, 38.5 ± 12% (n = 46 neurons) Y/C ratio increase in the soma, indicating robust detection of neuronal CaMKII activity (Figure 5a). Next, we tested if CaMKAR-CY can detect CaMKII activation induced by spontaneous activity. Cultured hippocampal neurons exhibit spontaneous synchronous firing, which is essential for proper network development and homeostasis. This intrinsic activity drives Ca^2+^ oscillations and the activation of signaling pathways, including CaMKII. In neurons co-expressing HaloCaMP1a^44^ to simultaneously visualize Ca^2+^ dynamics, we detected a 2.4 ± 1.5% (n = 20 neurons) CaMKAR-CY response in the soma following individual firing events (Figure 5b). This small but reproducible increase suggests a transient activation of CaMKII in response to spontaneous neuronal firing. In contrast, no Y/C ratio increase was observed with CaMKAR-CY (T/A) (Supplementary Fig. 10a).

**Figure 5.**
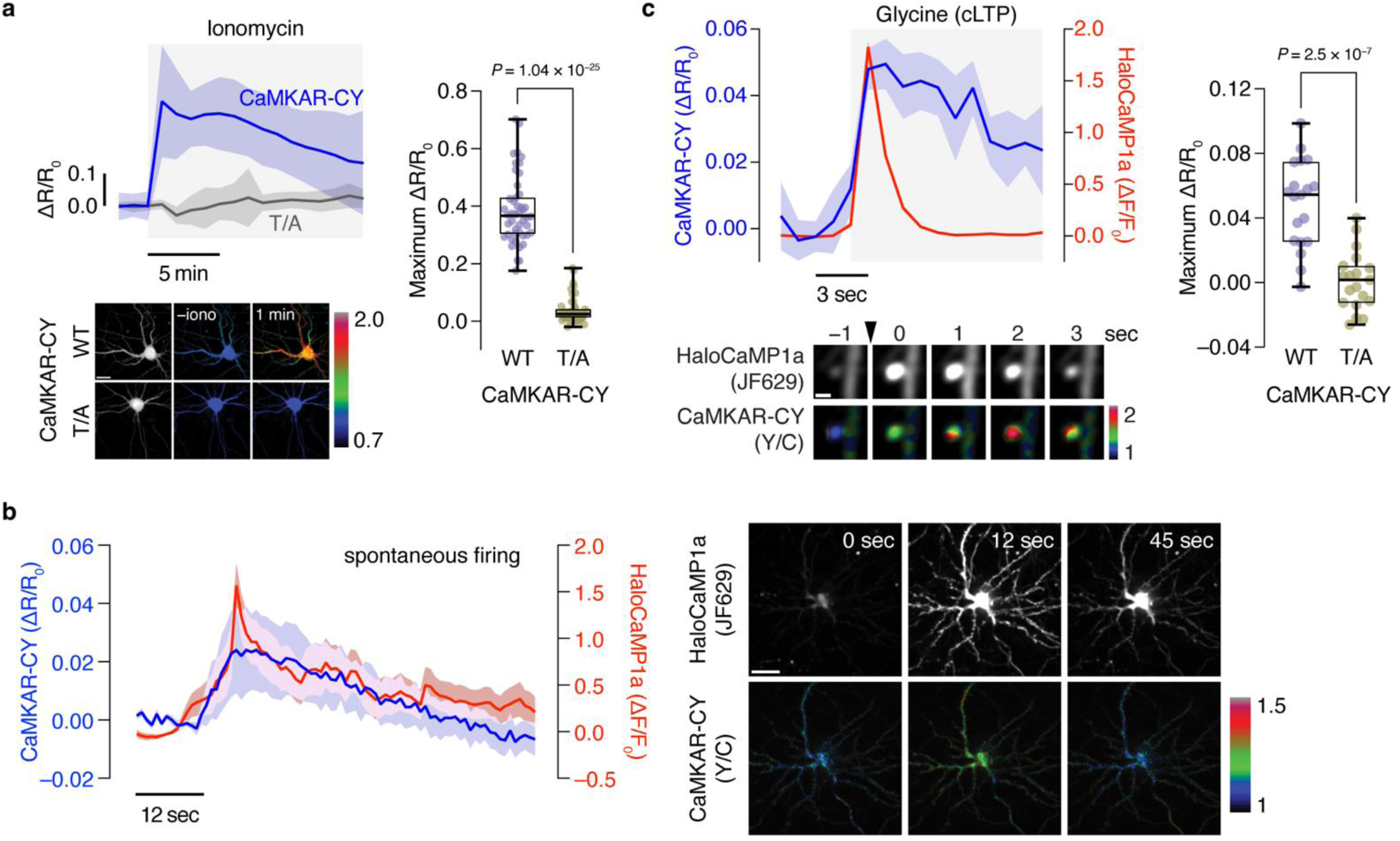
Sensitive detection of neuronal CaMKII activity. **a**, Top left, average timecourses of CaMKAR-CY and CaMKAR-CY (T/A) ratio change in hippocampal neurons treated with 5 μM Iono. n=46 cells from 3 independent experiments. Bottom left, representative images showing the Y/C ratio (pseudocolored) before and after Iono stimulation for CaMKAR-CY (top) and CaMKAR-CY (T/A) (bottom). Right, summary of the maximum Iono-stimulated CaMKAR-CY responses. **b**, Left, average responses from HaloCaMP1a (red) and CaMKAR-CY (Y/C, blue) aligned to Ca^2+^ peaks from spontaenous firing events. n=20 cells from 3 independent experiments. Right, representative images of a spontaneous firing event (HaloCaMP1a, upper) and corresponding CaMKAR-CY ratio images (lower). **c**, Top left, average responses from HaloCaMP1a (red) and CaMKAR-CY (blue) aligned to Ca^2+^ peaks detected following glycine addition to induce cLTP. Bottom left, representative images of an identified dendritic spine showing a Ca^2+^ transient event (upper) following cLTP induction (arrowhead) and corresponding CaMKAR-CY ratio images (lower). Right, summary of the maximum responses for CaMKAR-CY and CaMKAR-CY (T/A) following Ca^2+^ peaks. n=19 cells from 3 independent experiments each. Solid lines in timecourses indicate means; shaded areas, s.e.m. Box-and-whisker plots show the median, quartiles, min, and max. Unpaired, two-tailed Student’s t-test with (**a**) or without (**c**) Welch’s correction. Scale bars, 20 μm (**a**,**b**) and 2 μm (**c**). See also Supplementary Fig. 10.

Ca^2+^ influx through N-methyl-D-aspartate (NMDA) receptors is a critical event that activates multiple downstream signaling pathways during synaptic plasticity, including CaMKII^3^. To further explore the spatiotemporal interaction between Ca^2+^ entry and CaMKII activity during synaptic plasticity, we used a widely adopted chemical long-term potentiation (LTP) protocol^45^, which induces robust synaptic strengthening by mimicking physiological patterns of synaptic activity. As above, we simultaneously visualized Ca^2+^ dynamics and CaMKII activity. Upon glycine stimulation, which activates NMDARs and induces Ca^2+^ transients in spines, we observed a 183 ± 5% (n = 19 neurons) increase in HaloCaMP1a fluorescence intensity (Figure 5c). Immediately following individual postsynaptic Ca^2+^ transients, we also observed local CaMKII activity increases, as indicated by the CaMKAR-CY emission ratio (ΔR/R_0_ = 4.8 ± 0.6%). These activity increases subsequently decayed over tens of seconds, highlighting the transient nature of CaMKII activation during plasticity events. In contrast, CaMKAR-CY (T/A) showed no Y/C ratio (Supplementary Fig. 10b), further confirming CaMKAR-CY specificity.

CaMKII is directly recruited to dendritic spines through interactions with NMDA receptor subunits, and imaging of spine-localized CaMKII signaling is crucial for unravelling the spatiotemporal regulation of synaptic plasticity^3^. However, direct visualization of CaMKII activity dynamics in dendritic spines has yet to be achieved, as previous efforts have exclusively relied on monitoring CaMKII conformational changes as an indirect proxy. We therefore set out to image CaMKII activity in single dendritic spines via 2-photon fluorescence lifetime imaging (2pFLIM), leveraging the inherent FLIM-compatibility of FRET-based biosensors^46,47^. We optimized CaMKAR-CY for FLIM compatibility by replacing the original Cerulean donor with EGFP to allow more red-shifted excitation and the cpVenus[E172] acceptor with cpsREACh, a circularly permuted dark YFP^48,49^ to minimize spectral cross-talk, yielding CaMKAR-FLIM (Fig. 6a). Both CaMKAR-CY and CaMKAR-FLIM exhibited robust fluorescence lifetime changes in response to Iono stimulation (Fig. 6a-b and Supplementary Fig. 11a), substantially outperforming both FRESCA and FRESCA-2sRet, a FLIM-optimized FRESCA containing 2sREAChet^50^ and EGFP. Given it’s more favorable spectral characteristics, CaMKAR-FLIM was used for the subsequent experiments.

**Figure 6.**
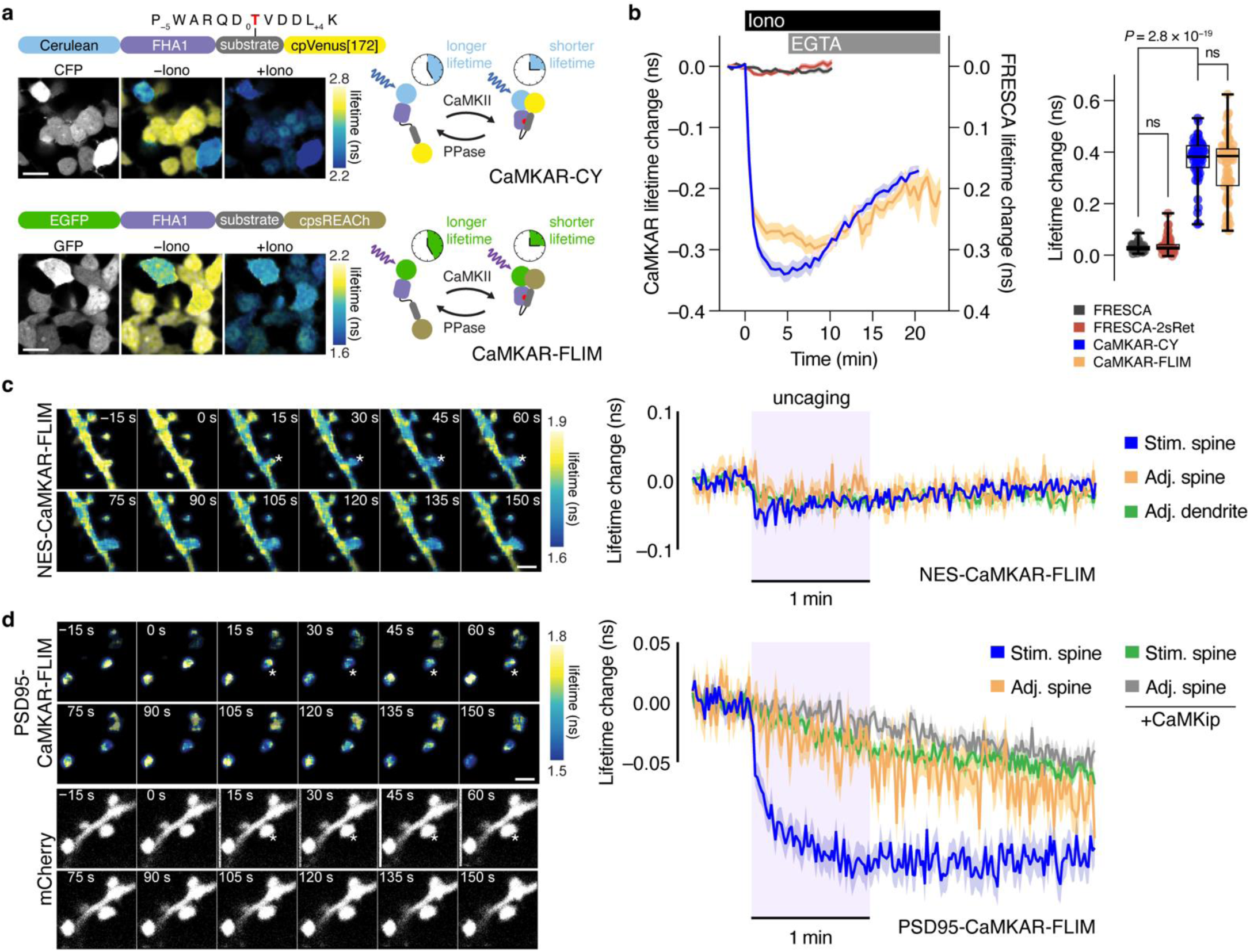
Lifetime imaging of CaMKII activity in hippocampal CA1 synapses. **a**, Representative fluorescence intensity and lifetime (pseudocolor) images of HEK293 cells expressing CaMKAR-CY and CaMKAR-FLIM before and after Iono application. Scale bars, 20 μm. **b**, Left, averaged timecourses of fluorescence lifetime change of CaMKAR-CY (n=55 cells), CaMKAR-FLIM (n=67 cells), FRESCA (n=41 cells), or FRESCA-2sRet (n=73 cells) in HEK cells treated with Iono (12 μM) followed by EGTA (10 mM). Vertical axis of FRESCA sensors inverted for comparison. Right, maximum fluorescence lifetime changes, plotted as absolute values for comparison. Box-and-whisker plots show median, quartiles, min, and max. ns, *P* > 0.999; Kruskal-Wallis test followed by Dunn’s multiple-comparisons test. **c-d**, Left, representative fluorescence lifetime images of dendritic spines during sLTP induction via glutamate uncaging in CA1 pyramidal neurons expressing (**c**) NES-CaMKAR-FLIM or (**d**) PSD95-CaMKAR-FLIM (upper images) and mCherry as a morphology marker (lower images) in organotypic slice culture. Asterisks indicate the location of glutamate uncaging. Scale bars, 2 μm. Right, average timecourses of fluorescence lifetime change of (**c**) NES-CaMKAR-FLIM and (**d**) PSD95-CaMKAR-FLIM in the stimulated (Stim) spine, adjacent (Adj) spine (within ∼7 μm of Stim spine), and adjacent dendrite (within ∼4 μm of Stim spine). n (spines/neurons)=42/4 (**c**) and 45/10 (**d**). Responses from Stim and Adj spines in slices expressing PSD95-CaMKAR-FLIM plus mCherry-CaMKip (n=94/19) are also shown in **d**. Solid lines and shaded areas in timecourses indicate mean and s.e.m., respectively. See also Supplementary Fig. 11.

We then biolistically transfected organotypic hippocampal slices with CaMKAR-FLIM and performed 2pFLIM in the dendrites of CA1 pyramidal neurons (Fig. 6c-d), using 2p glutamate uncaging in the absence of extracellular Mg^2+51^ to induce structural long-term potentiation (sLTP) in single spines. Compared with the CaMKII activation sensor 2dV-Camui^52^, which showed a large lifetime signal confined to the stimulated spine^52–54^ (Supplementary Fig. 11b), CaMKAR-FLIM showed a more muted response that spread across both stimulated and adjacent spines, also reaching the stimulated dendrite (Fig. 6c). We hypothesized that this spatially broad response pattern was likely due to sensor diffusion^50^ and thus fused CaMKAR-FLIM to PSD-95, one of the most abundant proteins in the postsynaptic density^55^. As expected, PSD95-CaMKAR-FLIM was concentrated into puncta within spines and showed a large lifetime response, which was confined to the stimulated spines without invading surrounding spines (Fig. 6d). Notably, the CaMKAR-FLIM response was more prolonged compared to that of 2dV-Camui, which exhibited decay time constants of 5 s (62%) and 220 s (38%). This reflects a key difference in the sensor design – Camui measures kinase activation, whereas CaMKAR detects kinase activity by measuring substrate phosphorylation. Importantly, no CaMKAR-FLIM response was detected in slices co-expressing CaMKip-mCherry (Fig. 6d), confirming CaMKII specificity. Using CaMKAR-FLIM, we are thus able to achieve direct, live-cell visualization of endogenous CaMKII activity in dendritic spines. Our findings represent a major advance over past efforts reliant on indirect conformational readouts using overexpressed kinase and showcase the versatility and broad applicability of these tools.

Overall, these results clearly demonstrate the utility of FRET-based CaMKARs for specific and rapid monitoring of endogenous CaMKII activity across diverse biological contexts and imaging modalities. The ability to track spatiotemporal CaMKII activity dynamics is crucial for unraveling the molecular mechanisms underlying key biological processes, and these tools will be instrumental in furthering our understanding of CaMKII signaling.

## DISCUSSION

In this study, we develop a suite of sensitive FRET-based CaMKII kinase activity reporters capable of robustly detecting endogenous CaMKII activity in various biological systems. FRESCA2 shows a 30-fold higher dynamic range over our previous CaMKII sensor FRESCA^13^, primarily driven by exchanging the original FHA2 domain for FHA1 (Fig. 1 and Supplementary Fig. 1). We also generated CaMKAR-CY, containing a redesigned CaMKII substrate peptide with increased sensitivity and selectivity, which showed an ∼130% maximum response upon CaMKII activation in HeLa cells and no response to PKD or other CaMK family kinases (Fig. 2). CaMKAR-CY dynamic range is nearly identical to that of a recently reported excitation-ratiometric CaMKAR based on a single-FP design^27^ (Supplementary Fig. 6). However, we found that CaMKAR-CY responded significantly faster to Ca^2+^ elevation than excitation-ratiometric CaMKAR (Supplementary Fig. 6), showing that CaMKAR-CY is more sensitive. Importantly, FRESCA, FRESCA2, and CaMKAR-CY all showed minimally increased responses upon phosphatase inhibition (Supplementary Figs. 1 and 8), confirming that dynamic range was not limited by dephosphorylation and that performance improvements were exclusively due to enhanced kinase sensitivity.

Using CaMKAR-CY, we were able to visualize pacing-induced elevations in CaMKII activity in primary cardiomyocytes (Fig. 3), as well as spontaneous and cLTP-triggered CaMKII activity changes in dissociated hippocampal neurons (Fig. 5). FRESCA2 exhibited faster reversal than CaMKAR-CY, likely reflecting more efficient dephosphorylation. CaMKAR-CY nevertheless successfully enabled extended imaging of low-frequency CaMKII activity oscillations under fertilization-mimicking conditions in mouse oocytes (Fig. 4). Notably, CaMKAR-CY signal decay is likely limited by dephosphorylation rather than CaMKII inactivation kinetics, as Camui responses, which directly reflect CaMKII activation, decay rapidly under similar conditions^13^ {ref}. Future efforts will thus focus on modifying the CaMKAR-CY substrate or tuning FHA1-phosphosubstrate affinity to enhance dephosphorylation. We further constructed CaMKAR-FLIM to facilitate 2pFLIM imaging in hippocampal slices. Having the ability to detect endogenous CaMKII activity dynamics within single dendritic spines during structural LTP (Fig. 6) opens many possibilities for interrogating the role of CaMKII kinase activity in learning and memory, which has recently come under debate^56,57^. The immediate compatibility of FRET-based sensors with quantitative approaches such as FLIM is a distinct advantage over single-FP sensors, which typically require bespoke re-engineering^58,59^. Though titration of endogenous activity is always a concern when imaging substrate-based probes, these concerns can be addressed by ensuring that sensor performance is independent of fluorescence intensity (i.e., expression level), reducing sensor DNA amounts, and imaging less-bright cells. Our versatile and broadly applicable FRET-based CaMKARs are thus a powerful addition to the CaMKII sensor toolkit.

Genetically encoded fluorescent kinase activity reporters (KARs) have transformed the study of kinase signaling by enabling direct, real-time visualization of kinase activity dynamics with spatiotemporal precision^9^. Yet KAR development is often slowed by a lack of rational, systematic engineering approaches, leading many kinases to be underserved by this technology. Designing substrate peptides that are robustly and selectively phosphorylated by the kinase of interest remains a particular challenge. Sequences derived from endogenous kinase substrates often provide a useful starting point but can be misleading in terms of their selectivity. Indeed, the endogenously derived substrate sequences used in various ERK sensors were recently shown to sense Cdk1 phosphorylation^60^. We similarly observed weak phosphorylation of FRESCA2 by PKD in some cells, consistent with the reported *in vitro* preferences of syntide-2^15,21^. However, recent systematic investigations of the substrate preferences for nearly all Ser/Thr^26^ and Tyr^61^ kinases offer a promising solution. Our approach of using Ser/Thr kinome atlas data to quantitatively predict selectivity-driving mutations, and thus re-engineer our CaMKII substrate peptide (Fig. 2), can serve as a general strategy for rapidly improving the selectivity and performance of KARs. Development of computational pipelines leveraging these data may ultimately help automate substrate design to enable dramatically accelerated KAR engineering.

In summary, we have developed a suite of FRET-based CaMKII activity reporters based on a generalizable rational design strategy. FRESCA2, CaMKAR-CY, and CaMKAR-FLIM comprise a powerful toolkit for probing the spatiotemporal dynamics of endogenous CaMKII signaling in living cells and offer a robust proof of concept for future efforts to expand kinase engineering and illuminate previously unmapped aspects of kinase signaling biology.

## ONLINE METHODS

### Plasmids

FRESCA^13^ and 2dV-Camui^52^ have been described previously. The FRET-based PKD sensor DKAR^22^ was kindly provided by Alexandra Newton (UC San Diego). The red Ca^2+^ sensor RCaMP1d^62^ was a gift of Loren Looger (UC San Diego), and the far-red Ca^2+^ sensor NIR-GECO2G^19^ was a gift of Robert Campbell (University of Tokyo). pAAV-synapsin-HaloCaMP1a-EGFP^44^ was a gift from Eric Schreiter (Addgene plasmid #138327) and was subcloned into pCAGGS using standard molecular cloning techniques. The dsDNA sequence encoding mouse CaMKIIN1 protein (UniProt: Q6QWF9) was synthesized by Genewiz (Azenta Life Sciences) and cloned into the pGEMHE vector using Gibson assembly. The resulting construct was confirmed by whole plasmid sequencing.

To construct FRESCA2, the FHA1 domain of AKAR4^17^ was PCR-amplified to incorporate the syntide-Thr substrate sequence (i.e., S1) from FRESCA at the FHA1 C-terminus, followed by insertion into *Sph*I/*Sac*I-digested AKAR4 in pRSETb via Gibson assembly^63^. The construct was then subcloned into pcDNA3 via *Bam*HI/*Eco*RI digestion. Substrate variants S2-S4 were subsequently generated via Gibson assembly following PCR amplification of pcDNA3-FRESCA2 using primer pairs encoding the corresponding mutations in the substrate region. Variant S4 was designated CaMKAR-CY. Non-phosphorylatable, T/A negative-control sensor mutants were cloned in a similar fashion. The CaMKip-mCherry inhibitor construct was generated by PCR-amplification of mCherry to add 5’ Kozak^64^ and AIP2^20^ sequences, followed by Gibson assembly into *Hind*III/*Eco*RI-digested pcDNA3.1(+).

To construct CaMKAR-FLIM, a fragment encoding the FHA1 domain and S4 substrate sequences from CaMKAR-CY was PCR-amplified to add an EGFP-overlapping region at the FHA1 N-terminus, and cpsREACh was PCR-amplified from tAKARα^65^ (gift of Haining Zhong, Addgene Plasmid #119913) to add S4 substrate- and pcDNA3.1-overlapping regions at the cpsREACh N- and C-termini, respectively. The resulting PCR fragments were inserted via Gibson assembly into a pcDNA3.1 backbone containing EGFP. NES-CaMKAR-FLIM was generated by inserting a PCR fragment encoding CaMKAR-FLIM into an *Age*I/*Eco*RI-digested pCI backbone containing an N-terminal NES (MLQNELALKLAGLDINKTG) via Gibson assembly. PSD95-CaMKAR-FLIM was then generated by inserting a PCR fragment encoding full-length PSD95 into *Not*I/*Age*I-digested NES-CaMKAR-FLIM via Gibson assembly. FRESCA-2sRet was generated by PCR-amplification of two sREAChet, FHA2, and syntide-EGFP-NES fragments and insertion into *NheI*/*Not*I-digested pCI vector via Gibson assembly. To construct CaMKIIα-IRES-CaM, a PCR fragment encoding full-length CaM was inserted into *Bst*XI/*Not*I-digested pIRES2-EGFP (Clonetech) via Gibson assembly to obtain pIRES2-CaM. An *Nhe*I/*Bam*HI-digested PCR fragment encoding full-length CaMKIIα was then ligated into *Nhe*I/*Bam*HI-digested pIRES2-CaM. The contruct was then subcloned behind a CAG promoter via *Nhe*I/*Not*I digestion. Gibson assembly reactions were performed using NEBuilder HiFi DNA Assembly kit (New Englad Biolabs). All plasmids were verified by Sanger sequencing or whole-plasmid sequencing. Biosensor sequences are shown in Supplementary Fig. 12.

### PSSM data analysis

Normalized and scaled PSSM data were downloaded from the Ser/Thr kinome atlas^26^. Matrices for the kinase of interest (KOI: CAMK2A) and kinase of detriment (KOD: PRKD1) were subjected to elementwise division to determine selective sites. Within the selectivity matrix, a value <1 represents a KOD-selective site, a value of 1 represents a site that is equally attractive or unattractive to both the KOI and KOD, and a value >1 represents a KOI-selective site. To ensure KOI sensitivity is equivalently weighed with selectivity, the KOI matrix was then linearly rescaled to match the range of the selectivity matrix and averaged with the selectivity matrix to obtain the final substrate matrix used for substrate peptide sequence design.

### Substrate peptide phosphorylation assays

*In vitro* kinase activity assays were performed similar to previous reports^66,67^. Varying concentrations of CaMKII substrate peptides (Genscript) were incubated in reaction buffer containing 105 mM Tris-HCl, pH 8.0, 150 mM KCl, 10 mM MgCl_2_, 2 mM ATP, 1 mM phosphoenolpyruvate (Alfa Aesar), 0.2 mM NADH (Sigma), 10 U/ml Pyruvate kinase (Sigma), 30 U/ml Lactate dehydrogenase (Millipore Sigma), and 5 nM CaMKIIα catalytic domain (residues 7-274) for 10 min at 30 °C. NADH fluorescence was measured at 450 nm at 20-s intervals using a Synergy H1 microplate reader (Biotek). Reaction rates were calculated by the maximum slope of NADH fluorescence over an 80-s period. Data were fitted to the Michaelis-Menten equation in GraphPad Prism 10 (GraphPad Software).

### Cell culture and transfection

#### Cell lines

HeLa cells were cultured in Dulbecco modified Eagle medium (DMEM; Gibco) containing 1 g L^−1^ glucose, 10% (v/v) fetal bovine serum (FBS, Sigma), and 1% (v/v) penicillin-streptomycin (Pen-Strep, Sigma-Aldrich) Cells were maintaining in a humidified incubator at 37 °C and 5% CO_2_. Cells were passaged every 3-4 d and routinely checked for mycoplasma contamination via regular DNA staining. For imaging experiments, cells were plated onto sterile 35-mm glass-bottomed dishes and grown for 24 h to reach 50-70% confluence. Transient transfection of biosensor-expressing plasmids was performed using calcium-phosphate precipitation. HeLa cells were given fresh culture medium 24 h after transfection and then incubated for an additional 6 h prior to imaging.

HEK-tsA201 (Sigma-Aldrich, St. Louis, US) were cultured in T75 flasks at 37 °C, 5 % CO2 in complete DMEM with 4.5 g L^−1^ glucose (Sigma-Aldrich). Culture media was supplemented with 10 % (v/v) fetal bovine serum (FBS; Capricorn, Ebsdorfergrund, Germany), 100 U mL^−1^ penicillin, 100 mg mL^−1^ streptomycin (Sigma-Aldrich), and 2 mM L-glutamine (Sigma-Aldrich). For single-cell fluorescence microscopy experiments, HEK-tsA201 cells were seeded on 25 mm glass coverslips (Glaswarenfabrik Karl Hecht, Sondheim vor der Rhön, Germany) in 6-well plates at a density of 2.5 × 10^5^ cells/well in 2 mL culture medium. The coverslips were coated before seeding with Poly-D-Lysine (PDL; Sigma-Aldrich; 25 µg mL^−1^) for 30 min at room temperature and washed twice with Dulbecco’s Phosphate Buffered Saline (PBS; Sigma-Aldrich). Cells were transfected 24 h after seeding using Lipofectamine™ 2000 transfection reagent (Thermo Fisher Scientific, Waltham, US). To transfect cells on one coverslip, 1.5 µg cDNA was premixed with 150 µL Opti-MEM (Thermo Fisher Scientific). In a second tube, 3.75 µL Lipofectamine™ 2000 transfection reagent was premixed with 150 µL Opti-MEM and, after 5 min incubation at room temperature (RT), both solutions were combined. After 20 min incubation at RT, the transfection mixture was added dropwise to the well.

#### Primary cells

##### Cardiac Myocytes

All rabbit handling and laboratory procedures were performed in accordance with the approved protocols of the Institutional Animal Care and Use Committee at University of California, Davis (#23175) conforming to the Guide for the Care and Use of Laboratory Animals published by the US National Institutes of Health (8^th^ edition, 2011).

Cardiac ventricular myocytes were isolated from male New Zealand White rabbits (3-4 months old, 2.5-3 kg), as previously described^68,69^. Briefly, rabbits were injected with heparin (400 U kg^−1^ body weight) before induction of general anesthesia by one-time intravenous injection of propofol (Rapanofal®, Ivaoes Animal Health, Miami, FL, USA; 10 mg/kg body weight) followed by isoflurane inhalation (2-5% in 100% O_2_). Deep surgical anesthesia was confirmed by abolishment of pain reflexes prior to euthanasia by surgical excision of the heart. Immediately after excision, the heart was rinsed in cold nominally Ca^2+^-free perfusion solution containing (in mM): 135 NaCl, 5.31 KCl, 1 MgCl_2_, 0.33 sodium phosphate monobasic, 2 Na pyruvate, 10 HEPES Na salt, 10 HEPES free acid (VWR), 5.5 glucose; 0.04% insulin, 0.5% Penicillin-Streptomycin, heparin (5 U ml^−1^); pH 7.4, Trizma® base. The aorta was cannulated and retrograde-perfused with perfusion solution (20 µM [Ca^2+^], heparin 4 U mL^−1^, 37 °C, gassed with 100% O_2_) on a constant-flow Langendorff apparatus and digested with Collagenase Type II (Worthington Biochemical Corp., Lakewood, NJ) and Protease Type XIV. Myocytes were mechanically dispersed, filtered through a nylon mesh, and allowed to sediment for ∼10 min in solution containing 2% bovine serum albumin (BSA) to quench enzymatic activity. The sedimentation was repeated two times while increasing [Ca^2+^] from 10 to 25 µM and decreasing BSA in 1% increments. Isolated myocytes were kept at room temperature until use.

Isolated myocytes were plated 1.2 × 10^4^ cells per well on laminin-coated µ-Slide 8 Well high chamber slides (ibidi, polymer #1.5) and cultured for 24-48 h at 37 °C in M199 medium (Gibco, Thermo Fisher Scientific, Waltham, MA) supplemented with insulin-transferrin-selenium (Gibco, Thermo Fisher Scientific, Waltham, MA), 1% Penicillin-Streptomycin. Myocytes were co-infected with purified CaMKAR-CY adenovirus (MOI: 3.3; 1.0 × 10^11^ particles mL^−1^) and purified calmodulin (CaM) adenovirus (MOI: 3.0; 8.0 × 10^9^ particles mL^−1^; a gift from David Yue, The Johns Hopkins University, Baltimore, MD) to compensate for a decrease in endogenous CaM expression in culture^70^ and to facilitate CaMKII activation and response to field-stimulation.

The adenoviral vector for CaMKAR-CY was generated per the AdEasy XL Adenoviral Vector System (Agilent, Santa Clara, CA). In brief, CaMKAR-CY cDNA was subcloned into pShuttle-CMV plasmid and verified by restriction analysis and sequencing. Adenoviral DNA was generated by homologous recombination and then transfected into HEK293 cells for amplification. The amplified adenovirus was then purified by CsCl gradient and dialyzed into solution composed of (in mM): 1 Tris-HCl, 1 MgCl_2_; 5% glycerol; pH 7.8. Titration was determined spectrophotometrically (OD_260_). All reagents were purchased from Sigma-Aldrich unless indicated otherwise.

##### Mouse Eggs

Mouse eggs were collected from B6D2F1 mice bred from B6 and DFA2 parental lines (Jackson Laboratory) at the UMass animal facility. The University of Massachusetts Institutional Animal Care and Use Committee (IACUC) approved all animal experiments and protocols. Mature oocytes (MII eggs) were extracted from the ampulla of female mice aged 6 to 8 weeks. Females were induced to undergo superovulation via intraperitoneal injection of 5 IU of pregnant mare serum gonadotropin (PMSG, Sigma, St. Louis, MO) followed by 5 IU of human chorionic gonadotropin (hCG, Sigma) at 48-h intervals. Cumulus-oocyte complexes (COCs) were collected 14-16 h after hCG injection by carefully tearing the ampulla with forceps and needles in TL-HEPES medium. The COCs were then treated with 0.26% (w/v) hyaluronidase (Sigma-Aldrich, H3506) at room temperature for 5 min to facilitate the removal of cumulus cells. After collection, oocytes were recovered in potassium simplex optimized medium (KSOM) until microinjection. KSOM dishes was covered with light mineral oil (Fisher Scientific, Hampton, NH), and then incubated in a humidified atmosphere with 5% CO_2_ at 37 °C.

Prior to *in vitro* transcription, FRESCA, FRESCA2, and CaMKAR-CY plasmids were linearized via *Xba*I digestion, and the CaMKIIN1 plasmid was linearized with *Asc*I. Transcription was performed using the T7 mMESSAGE MACHINE kit (Thermo Scientific, AM1344). Poly(A)-tails were added to mRNAs using a Poly(A) Tailing kit (Thermo Scientific, AM1350). mRNAs were then purified using Megaclear™ Transcription Clean-Up Kit (Thermo Scientific, AM1908), and 3 µL mRNA aliquots were stored at -80 °C. Purified mRNAs were denatured at 95°C centrifuged for 10 min at 4 °C 13000 rpm, and the top 2 μL was then used to prepare microdrops from which glass micropipettes were loaded by aspiration. mRNA (1 μg μL^−1^) was delivered into eggs by pneumatic pressure (PLI-100 pico injector, Harvard Apparatus, Cambridge, MA). Each egg received ∼5–10 pL, corresponding to ∼1–3% of the total intracellular volume. Injected MII eggs were incubated for in KSOM up to 5 h to allow for translation of the biosensor constructs.

##### Neurons

Rat hippocampal neurons were cultured as previously described^71^. Briefly, hippocampal neurons obtained from embryonic day 18 Sprague-Dawley rats were plated on poly-L-lysine-coated glass cover slips at a density of 150,000 cells/well in NM5 (neurobasal media (Gibco) supplemented with 2% B-27, 4 mM GlutaMax, 50 U/mL PenStrep, and 5% horse serum (Hyclone)). Culture media was changed the next day to NM0 (NM5 minus horse serum), and neurons were fed once a week with NM0. At 15-18 days in vitro, neurons were transfected with the indicated plasmids using Lipofectamine 2000 (Invitrogen). All experimental procedures involving animals were conducted according to the National Institutes of Health guidelines for animal research and were approved by the Animal Care and Use Committee at Johns Hopkins University School of Medicine.

##### Organotypic hippocampal slice cultures

All experimental procedures were approved and carried out in accordance with the regulations of the Max Planck Florida Institute for Neuroscience Animal Care and Use Committee, as per US National Institutes of Health guidelines. Organotypic cultured hippocampal slices were prepared from postnatal 5-9 days C57BL/6 mice, as described^52,72^. After 7-8 days in culture, CA1 pyramidal neurons were transfected via biolistic gene transfer^73^ (Bio-Rad) with gene bullets made from 1.6-μm gold beads (8 mg) coated with plasmids containing cDNA of 2dV-Camui (15 μg), CaMKAR-FLIM (15 μg), or PSD95-CaMKAR-FLIM (15 μg) and mCherry (10 μg). 2pFLIM imaging was performed 2-8 days after transfection.

### Time-lapse fluorescence imaging

#### HeLa cell imaging

Thapsigargin (Tg; Cayman Chemical, 10522), ionomycin (Iono; Calbiochem, 407950), Calyculin A (CalA; LC Labs, C-3987), Phorbol 12-myristate 13-acetate (PMA; LC Labs, P-1680), and CRT0066101 (CRT; Tocris, 4975) were prepared at 1000X stock concentration in DMSO (Fisher, BP231). Calcium chloride (CaCl_2_; Fisher, C79) was prepared at 2 M stock concentration in ultrapure water.

HeLa cells were washed twice with Ca^2+^-free imaging buffer (HBSS, Ca^2+^-free, supplemented with 20 mM HEPES, pH 7.4, 2 g L^−1^ glucose, and 100 μM EGTA) and subsequently imaged in Ca^2+^-free imaging buffer in the dark at 37 °C. Tg (1 μM), Iono (2 μM), CaCl_2_ (5 mM), CalA (10 nM), PMA (50 ng mL^−1^), and CRT (5 μM) were added as indicated. Cells were imaged on a Zeiss AxioObserver Z1 microscope (Carl Zeiss) equipped with a Definite Focus system (Carl Zeiss), a 40×/1.3 NA oil objective, and a Photometrics Evolve 512 EMCCD (Photometrics) and controlled by METAFLUOR 7.7 software (Molecular Devices). Dual cyan/yellow emission-ratio imaging was performed using a 420DF20 excitation filter, a 450DRLP dichroic mirror and two emission filters (475DF40 for CFP and 535DF25 for YFP). RFP intensities were imaged using a 555DF25 excitation filter, a 568DRLP dichroic mirror, and a 650DF52 emission filter. mIFP intensities were imaged using a 640DF30 excitation filter, a 660DLRP dichroic mirror, and a 700DF75 excitation filter. All filter sets were alternated using a Lambda 10-2 filter-changer (Sutter Instruments). All channels were acquired at 500 ms exposure time, except for YFP, which was 50 ms. EM gain was set at 50 for mIFP and 10 for all other channels. Images were acquired every 15 s.

Raw fluorescence images were analyzed using METAFLUOR 7.7 (Molecular Devices) software. Regions of interest (ROI) were drawn around biosensor-expressing cells, along with a cell-free region (background). Background-subtracted fluorescence intensities were calculated at each time point by subtracting the intensity of the background ROI from the intensities of each cell ROI and were then used to calculate emission ratios (yellow/cyan or cyan/yellow). All biosensor response timecourses were subsequently plotted as the normalized emission ratio or fluorescence intensity change with respect to time zero (e.g., ΔR/R_0_ or ΔF/F_0_), calculated as (R−R_0_)/R_0_ or (F−F_0_)/F_0_, where R and F are the emission ratio and fluorescence intensity value at a given time point, and R_0_ and F_0_ are the initial or emission ratio fluorescence intensity value at time zero, which was defined as the time point immediately preceding drug addition. Maximum ratio (ΔR/R) or intensity (ΔF/F) changes were calculated as (R_max_−R_min_)/R_min_ or (F_max_−F_min_)/F_min_, where R_max_ and R_min_ or F_max_ and F_min_ are the maximum and minimum emission ratio or intensity value recorded after stimulation, respectively. Maximum-normalized emission ratio changes (ΔR/R_max_) were calculated as (R_−drug_−R_+drug_)/(R_max_−R_0_), were R_−drug_ is the emission ratio value immediately prior to addition of the indicated drug, R_+drug_ is the maximum emission ratio recorded after drug addition, R_max_ is the overall maximum emission ratio recorded in the experiment, and R_0_ is the emission ratio value immediately preceding the 1^st^ drug addition. For CalA pre-treatment experiments, the residual response was evaluated using a sustained activity metric (SAM5), which was calculated as (R_drug,5 min_/R_max_), where R_drug, 5 min_ is the emission ratio value recorded 5 min after addition of the indicated stimulus and R_drug,max_ is the maximum emission ratio after stimulation. All graphs were plotted using GraphPad Prism 10 (GraphPad Software).

#### HEK-tsA201 cell imaging

HEK-tsA201 cells were imaged using an Olympus IX83 inverted microscope equipped with an oil immersion objective (UAPON40XO340-2 40X/1.35 oil) and an ORCA-Fusion C14440-20UP camera (Hamamatsu Photonics, Hamamatsu, Japan). A Spectra III-LCR-8S-A21 light engine (Lumencor, Beaverton, US) at 50 mW light intensity and an additional 20% transmission neutral-density filter (Qioptiq Photonics, Göttingen, Germany) were used for excitation with a 438/29 bandpass filter for CFP excitation, 511/16 for YFP direct excitation (CaMKAR-CY) and both 395/25 and 475/28 for cpGFP excitation (CaMKAR). In experiments with CaMKAR-CY, the emission light was split into two channels using an OPTOSPLIT II (Cairn research, Faversham, UK) with a 475/28 bandpass filter for CFP and a 542/27 bandpass filter for YFP. CaMKAR emission was collected with a 515/30 bandpass filter. Coverslips with transfected cells were transferred to imaging chambers (AttofluorTM, Thermo Fisher Scientific). Sequences of images were acquired with camera scan mode 2 and 2×2 camera binning. Acquisition control and live data view were managed by µManager^74^ in combination with custom Beanshell scripts. A constant superfusion of the cells with buffer flow and specific ligand addition were achieved by using a solenoid valve perfusion system with a 100-µm inner diameter manifold-tip (Octaflow II, ALA Scientific Instruments, Farmingdale, US) at a pressure of 350 mbar.

Image processing was performed with ImageJ2 (FIJI)^75^. The individual cells were marked as regions of interest (ROI). An area without cells was defined as ‘background’. Fluorescence intensity over time of all regions was extracted for each emission channel. The raw data were processed by subtracting the background fluorescence at every time point for all recorded emission channels. Data were analyzed using Excel (Microsoft). For the CaMKAR experiments, the ratio of R/R_0_ [%] was calculated after background correction as normalized differences between the basal emission ratio (Em515 with Ex475/ Em515 with Ex395) (R_0_) and the emission ratio at a given time point (R), by using the following equation: ΔR/R_0_[%]=(R−R_0_)/R_0_)×100. For FRET experiments, the acceptor emission was corrected for spectral bleed through (B) as Acceptor_emission_–B×Donor_emission_. The spectral bleed through (B) for the donor Cerulean was experimentally determined according to^76^. The FRET-ratios were calculated as the ratio of corrected acceptor emission (FRET) over donor emission. All data points were plotted as normalized differences between the basal FRET_0_ signal and the FRET signal at a given timepoint: ΔFRET[%]=(FRET−FRET_0_)/FRET_0_×100.

Plotting, curve fitting, and statistical analyses were performed using Prism 10.3 (GraphPad Software). The apparent on-rates (τ_on_) and the maximal efficacy (E_max_) were extracted by performing a plateau followed by one-phase association fitting (GraphPad model) as follows: F(t)=F_0_+(Plateau−F_0_)×(1–e^−K*(t-t^_0_^)^), if t<t_0_, where t is time [s], t_0_ the respective time point of ligand application, and Plateau−F_0_ (Span) the E_max_. From this, τ_on_ has been calculated as τ_on_=1/K.

#### Cardiac myocyte imaging

A Nikon Eclipse Ti confocal microscope (40X / 1.25 NA water immersion objective) controlled by NIS-Elements AR 4.30.02 software was used to acquire representative images (1024 x 1024 pixels; 0.203 µm/pixel) (Fig. 3a), fluorescence timecourses (Fig. 3b), and acceptor photobleach donor enhancement measurements (Supplementary Fig. 6a) of CaMKAR-CY expressed in rabbit ventricular myocytes. Donor CFP and acceptor YFP were excited using solid state laser illumination at 445 nm and 514 nm, respectively. Donor and acceptor emissions were collected at 460-500 nm (filter: 485/30) and 525-555 nm (filter: 540/30) detector ranges, respectively. Time series images (512 x 512 pixels) were acquired at a minimum rate of 2 Hz for timecourse traces. Donor enhancement measurements were acquired by photobleaching the acceptor with intense (100%) 514 nm laser exposure within circular regions of interest down to <5% of the initial acceptor fluorescence.

A Leica Falcon SP8 confocal microscope (40X / 1.30 NA oil immersion objective), equipped with variable hybrid detectors, controlled by Leica Application Suite X software was used to acquire fluorescence timecourses (Fig. 3c, Supplementary Fig. 6b, c). Donor CFP and acceptor YFP were excited using white-light femtosecond frequency-pulsed laser illumination at 440 nm and 514 nm, respectively. Donor and acceptor emissions were acquired at 450-500 nm and 525-600 nm HyD SMD hybrid detector ranges, respectively. Time series images (1024 x 1024 pixels) were acquired at a minimum rate of 2 Hz.

Rabbit ventricular myocytes expressing CaMKAR-CY were washed with nominally Ca^2+^-free Tyrode’s solution containing (in mM): 140 NaCl, 0 CaCl_2_, 1 MgCl_2_, 4 KCl, 5 HEPES Na salt, 5 HEPES free acid (VWR), 5.5 glucose; pH 7.4, NaOH. For CaMKII baseline activity measurements, myocytes were incubated in nominally Ca^2+^-free Tyrode’s solution (>30 minutes) to minimize CaMKAR-CY response before replacing the solution with Normal Tyrode’s solution containing 1.8 mM CaCl_2_. CaMKAR-CY response to physiological CaMKII activation was induced by pacing myocytes in Normal Tyrode’s solution (1.8 mM CaCl_2_) via field stimulation (pulse duration: 2 ms; amplitude: twice minimum action potential threshold) at 0.5 or 1 Hz, as indicated. β-adrenergic stimulation was induced by incubating myocytes for at least 3 minutes with 100 nM DL-isoproterenol hydrochloride (ISO). All experiments were performed at room temperature. All reagents were purchased from Sigma-Aldrich unless indicated otherwise.

CaMKAR-CY fluorescence time-courses were plotted (after cell-adjacent background subtraction for each wavelength) as average myocyte pixel fluorescence intensity (F), the emission ratio (R = F_DA_/F_DD_), or R normalized relative to that at the onset of steady state pacing (R/R_0_) or prior to Ca^2+^ addition (R/R_0Ca_). All plots were generated and statistical analyses were performed using GraphPad Prism 10 software.

#### Neuron imaging

Hippocampal neurons were imaged on a Zeiss spinning-disk confocal microscope equipped with a Yokogawa CSU-X1A 5000 spinning-disk unit, Photometrics Evolve EMCCD camera, six lasers (405/458/488/514/561/639 nm) and a Definite Focus system. FRET imaging of CaMKAR-CY was performed using the 458-nm laser line, a RTFT(457/514/647) dichroic mirror and 485/30 (CFP) 575/50 (YFP) emission filters. HaloCaMP1a labeled with Janelia Fluor 629 was imaged using the 639-nm laser, the RTFT(457/514/647) dichroic mirror and a 690/50 emission filter.

Neurons were imaged 2-4 days after transfection. Neurons were pre-incubated in 37 °C artificial cerebrospinal fluid (ACSF, 120 mM NaCl, 5 mM KCl, 2 mM CaCl_2_, 1 mM MgCl_2_, 10 mM D-glucose, and 10 mM HEPES, pH 7.4) for at least 60 min before being mounted on the microscope. HaloCaMP1a labeling (1 μM JF629-HTL) was performed for 1 hr in cell culture medium in a live-cell incubator, followed by two washes in ACSF at 37 °C. Coverslips with neurons were mounted on a heated, custom-built chamber without perfusion and imaged in ACSF warmed to 37 °C. For Iono stimulation, image stacks were acquired every 1 min. After 5 baseline images, 5 μM Iono was manually added to the chamber. Glycine stimulation was performed as previously described^71^. Briefly, neuron culture media were supplemented with 200 μM DL-AP5 1-2 days before experiments. On the day of experiments, neurons were preincubated in basal ACSF (ACSF supplemented with 200 μM DL-AP5, 1 μM TTX, 1 μM Strychnine, and 100 μM Picrotoxin) for at least 1 h before imaging. Images (fixed Z position) were acquired in streaming mode (∼1.1 Hz). Following 5-10 min of baseline recording, neurons were stimulated with glycine solution (ACSF without MgCl_2_, supplemented with 1 μM TTX, 1 μM Strychnine, 100 μM Picrotoxin, and 200 μM glycine). Image analysis was performed using ImageJ/Fiji as previously reported^71^.

#### Mouse egg imaging

Ionomycin calcium salt (Tocris; 1704), Calyculin A (CalA; LC Labs, C-3987), Phorbol 12-myristate 13-acetate (PMA; Tocris, 1201), Gö6983 (Tocris; 2285) and CRT0066101 (CRT; Tocris, 4975) were prepared at 1000X stock concentration in DMSO (Sigma-Aldrich, D8418). Dithiothreitol (DTT; Sigma-Aldrich, D0632) and strontium chloride hexahydrate (SrCl_2_•7H_2_O; Sigma-Aldrich, 255521) was prepared in ultrapure water at 1 M stock concentration.

Unless otherwise stated, all oocytes were monitored in a Ca^2+^ free HEPES-buffered Tyrode’s lactate solution (TL-HEPES) containing 5% heat-treated fetal calf serum (FCS; Gibco/ThermoFisher; Waltham, MA). For simultaneous monitoring of CaMKII and Ca^2+^ dynamics, eggs injected with biosensor mRNA were loaded with 2.5 μM Rhod-2AM (Invitrogen, R1244) and 0.02% pluronic acid (Invitrogen, P3000MP) for 30 min at room temperature approximately 5-6 h post-injection. Eggs were immobilized on glass-bottom dishes (MatTek Corp., Ashland, MA) by placing them in protein-free media, which caused them to adhere to the glass, and were then positioned on the stage of an inverted microscope. Imaging was conducted on a Nikon Eclipse TE 300 inverted epifluorescence microscope (Analis Ghent, Belgium) equipped with a 20X objective and a cooled Photometrics SenSys CCD camera (Roper Scientific, Tucson, AZ) controlled by NIS-elements software (Nikon). FRET and Rhod-2 imaging was performed using a three-color dichroic beamsplitter (89007 ET, Chroma) with YFP, CFP, and Rhod fluorescence collected using 430/470 nm, 500/535 nm, and 555/600 nm excitation and emission filters (Chroma Technology, Rockingham, VT), respectively. CFP, YFP, and Rhod-2 intensities were acquired at 900 ms exposure every 20 s. Raw fluorescence intensities were exported to Excel (Microsoft) from Nikon Elements software. Normalized FRET emission ratios (yellow/cyan or cyan/yellow) and Rhod-2 fluorescence intensities were calculated by dividing the raw ratio (R) or fluorescence intensity (F) at each time point by the average baseline value from the first 2 min of monitoring (e.g., R/R_baseline_ or F/F_baseline_). Oocyte data were also analyzed using custom Python scripts to identify peak locations, determine “start” and “end” boundaries, and then calculate amplitude, peak duration, and area under the curve. All graphs were plotted in GraphPad Prism 10 (GraphPad Software).

#### Two-photon fluorescence lifetime imaging

Fluorescence lifetimes were measured on a custom-built 2pFLIM system^52,77^ equipped with a Ti:Sapphire laser (Spectra-Physics, InSightX3) tuned to a wavelength of 920 nm for CaMKAR-FLIM and 850 nm for CaMKAR-CY. The laser power was modulated using a Pockels cell to 1.6-2.5 mW under the objective lens, and the laser position was scanned using XY galvo with 3-mm mirrors (Cambridge Technology, Model 6215H). Two-photon glutamate uncaging was performed with another Ti:Sapphire laser (Coherent, Chameleon) tuned to a wavelength of 720 nm and a power of 4.2-7.3 mW under the objective. A train of uncaging pulses was given with 30 pulses, 8-10-ms pulse width, and 0.5 Hz. Experiments were conducted in Mg^2+^-free ACSF (127 mM NaCl, 2.5 mM KCl, 4 mM CaCl_2_, 25 mM NaHCO_3_, 1.25 mM NaH_2_PO_4_ and 25 mM glucose) containing 1 μM TTX and 4 mM MNI-caged-L-glutamate (Tocris) aerated with 95% O_2_ and 5% CO_2_. The fluorescence was collected with a water-immersion objective (Olympus, LUMPlanFLN60× NA 1.0 W), divided by a dichroic mirror (565 nm) and detected with two separate photoelectron multiplier tubes (Hamamatsu Photonics, H7422-P40) placed after emission filters (Chroma, 520/60-2p for GFP or 470/30 for CFP, and 620/60-2p for mCherry).

Fluorescence lifetime imaging was obtained using a time-correlated single-photon counting board (Pico-Quant, Time-Harp 260) controlled with custom software (FLIMage)^52^. 2pFLIM images were taken of a 9.3×9.5 μm area at 128×128 pixels at 256 ms/frame with 4-frame average off-line. Experiments were performed at room temperature (23 ± 1 °C). Fluorescence lifetime imaging of HEK293 cells was performed in a HEPES-buffered ACF (20 mM HEPES pH 7.3, 130 mM NaCl, 2 mM NaHCO_3_, 25 mM D-glucose, 2.5 mM KCl and 1.25 mM NaH_2_PO_4_) with 2 mM CaCl_2_ and 2 mM MgCl_2_ by 2pFLIM as described above. Cells were stimulated with bath application of 12 μM ionomycin (Cell Signaling), followed by10 mM EGTA. ROIs for cells were generated by Cellpose^78^ for lifetime analysis.

### Statistics and reproducibility

All experiments were independently repeated as noted in the figure legends. All replication attempts were successful. Statistical analyses were performed using GraphPad Prism 10 (GraphPad Software). For Gaussian data, pairwise comparisons were performed using Student’s t-test or Welch’s unequal variance t-test, and comparisons among three or more groups were performed using ordinary one-way or two-way analysis of variance (ANOVA) followed by multiple comparisons tests as indicated in the figure legends. Non-Gaussian data were analyzed using the Mann-Whitney U test or Wilcoxon matched-pairs signed rank test for pairwise comparisons or the Kruskal-Wallis test followed by Dunn’s multiple comparisons test for analyses of three or more groups. Statistical significance was set at P < 0.05. Unless otherwise noted, averaged live-cell imaging timecourses depict mean±s.d.; box-and-whisker plots depict the median, quartiles, min, and max, as indicated in the figure legends.

## Data availability

Plasmids generated in this study will be made available through Addgene (www.addgene.org/Jin_Zhang). Source data are provided with the manuscript.

## Code availability

Custom ImageJ macros used to analyze neuronal imaging data and custom Jupyter Notebook Python codes used for PSSM analysis and for analyzing cardiac myocyte data are available upon reasonable request. Custom Python code used to analyze oocyte data (https://github.com/amanguliani/SensorAnalysisCode/blob/main/main.py) and FLIMage software for controlling FLIM hardware and analysing FLIM data are available on GitHub (https://github.com/ryoheiyasuda/flimage_public.git).

## ACKNOWLDEGEMENTS

The authors thank Alexandra Newton (UC San Diego), Loren Looger (UC San Diego), Eric Schreiter (Janelia Research Campus), Robert Campbell (U of Tokyo), and Haining Zhong (OHSU) for providing plasmids. We are also grateful to Maya Kunkel for helpful insights. We thank Richard C. Johnson, Yuan (Daisy) Zhou, Kelly South, and Anant Jain for help with subcloning, and Luke Lavis (Janelia Research Campus) for providing the Janelia Fluor 629 HaloTag ligand. We also thank the UC Davis Cardiovascular Research Institute Cell and Molecular Biology Core for help generating the recombinant CaMKAR-CY adenovirus, as well as Shannon Gilhooly and Anastasia Krajnovic (UC Davis) for help with cardiac myocyte isolation and culturing. This work was supported by NIH grants R35 CA197622 and R01 CA262815 (to J.Z.), R35 GM145376 (to M.M.S), R35 NS116804, R01 MH080047, and U01 NS128655 (to R.Y.), R01 HD092499 and R21 HD110734 (to R.A.F.), R01 HL142282 and R01 HL092097 (to D.M.B.), R35 CA197588, P01 CA120964, and P01 CA117969 (to L.C.C.); NIH BRAIN Initiative grant RF1 MH126707 (to R.L.H. and J.Z.); by the Deutsche Forschungsgemeinschaft (German Research Foundation) through SFB1423 (project number 421152132 - subprojects C05 (to A.B.) and C08 (to A.B. and M.J.L.)); a fellowship from the University of Massachusetts NIH Biotechnology Traineeship T32GM135096-01 (to N.A.T.); NSF Graduate Research Fellowship DGE-2038238 (any opinions, findings, and conclusions or recommendations expressed in this material are those of the author(s) and do not necessarily reflect the views of the NSF), AHA Predoctoral Fellowship 24PRE1186687, and NIH F31 Predoctoral Fellowship 1F31HL176051-01 (to A.C.L.); the UCSD Interfaces Graduate Training Program NIH/NIBIB T32 Training in Multi-scale Analysis of Biological Structures and Function Training Grant 5T32EB009380-15 (to S.A.S.); and funding from the Claudia Adams Barr Program for Cancer Research (to J.L.J.).

## AUTHOR CONTRIBUTIONS

S.M. and J.Z. conceived of the project. A.C.L. carried out computational analyses of raw kinome atlas scoring data provided by J.L.J., T.M.Y., and L.C.C.; S.M. designed CaMKII substrate sequences; S.M., S.A.S., and Y.N. generated biosensor constructs and performed live-cell imaging in HeLa and HEK293 cells; P.L. performed live-cell imaging in HEK-tsA201 cells; N.A.T. and O.C.K. performed live-cell imaging in mouse eggs; K.A. performed FLIM-FRET imaging in organotypic hippocampal slices; C.Y.K. performed live-cell imaging in adult cardiomyocytes using adenovirus generated by J.L.M.; B.L. and S.S.D. performed live-cell imaging in cultured hippocampal neurons; O.C.K. performed *in vitro* phosphorylation assays; S.M., A.B., M.J.L., L.C.C., D.M.B., R.Y., R.F., R.L.H., M.M.S., and J.Z. supervised the project and coordinated experiments; S.M., N.A.T., K.A., C.Y.K, B.L., S.S.D. P.L., and O.C.K. analyzed the data; and S.M., N.A.T., K.A., C.Y.K., B.L., P.L., A.B., D.M.B., R.Y., R.L.H., M.M.S., and J.Z. wrote the manuscript.

