## Supplementary Information for "Dynamic visualization of physiological CaMKII activity using sensitive FRET biosensors"

### Contents

**Supplementary Figures**

**Supplementary Table**

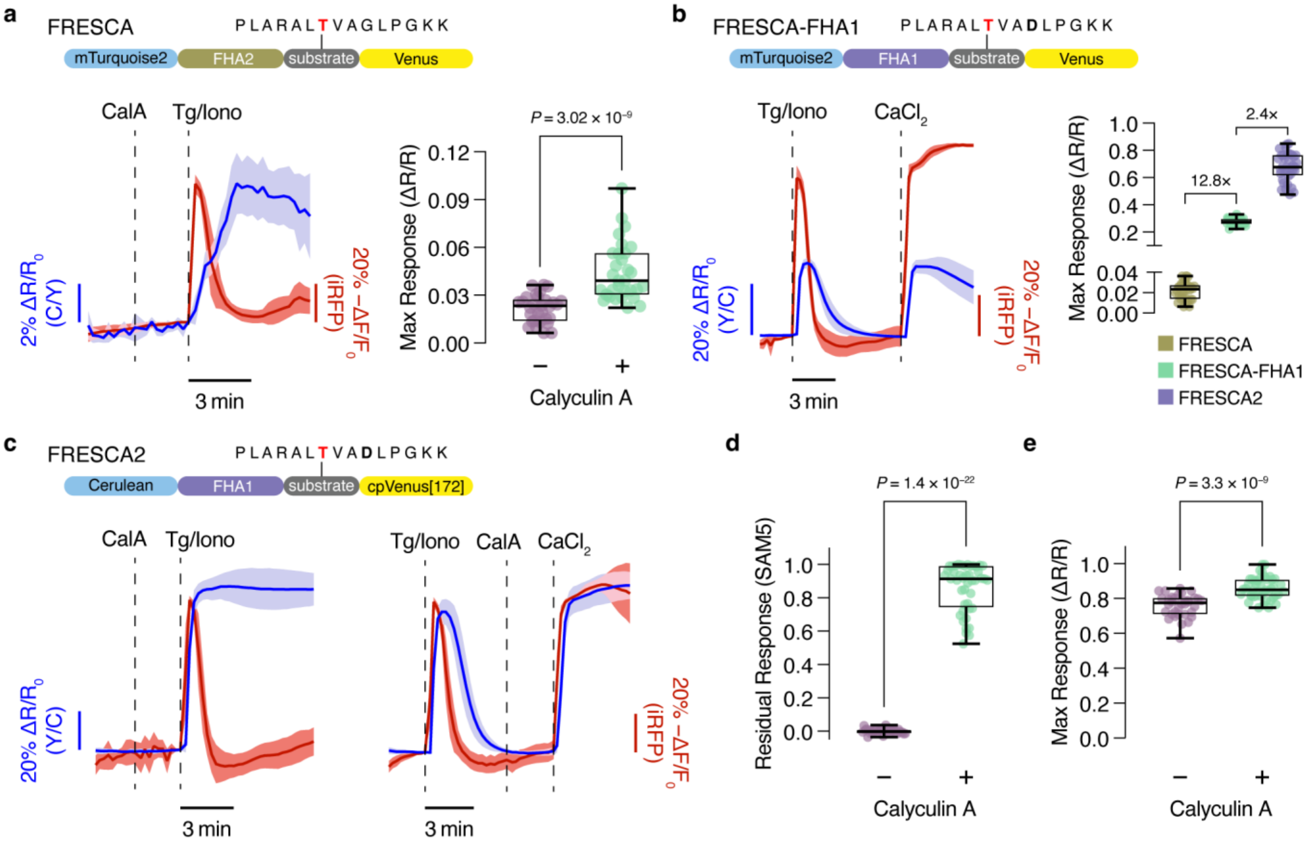


**Supplementary Figure 1 | Characterizing and improving FRESCA performance. a**, Phosphatase inhibition slightly increases FRESCA2 response. Left, representative averaged timecourse of FRESCA ratio (blue) and NIR-GECO2G intensity (dark red) responses in HeLa cells treated with 25 nM calyculin A (CalA) before Tg/Iono (n=7 cells). Data representative of 4 independent experiments. Right, summary of maximum FRESCA responses without (−; n=30 cells) or with (+; n=31 cells) CalA addition prior to Tg/Iono stimulation. Data are from 4 independent experiments. **b**, FHA1 domain insertion alone strongly enhances FRESCA performance. Right, representative averaged timecourse of FRESCA-FHA1 ratio (blue) and NIR-GECO2G intensity (dark red) responses in HeLa cells treated with Tg/Iono and $CaCl_2$ (n=10 cells). Data representative of 3 independent experiments. Right, summary of maximum Ca2+-stimulated responses from FRESCA (n=30 cells), FRESCA-FHA1 (n=35 cells) and

FRESCA2 (n=36 cells). Data from 3 independent experiments each. **c**, Phosphatase inhibition blocks reversal of FRESCA2 response. Representative averaged timecourses of FRESCA2 ratio (blue) and NIR-GECO2G intensity (dark red) responses in HeLa cells treated with 25 nM calyculin A (CalA) either before Tg/Iono (left, n=12 cells) or between Tg/Iono and CaCl2 (right, n=13 cells) stimulation. Data representative of 3 independent experiments each. **d** and **e**, Summary of (**d**) residual and (**e**) maximum FRESCA2 response without (−; n=34 cells) or with (+; n=44 cells) CalA addition prior to Tg/Iono stimulation. Data are from 3 independent experiments each. Solid lines in timecourse indicate the mean; shaded areas, s.d. Sensor schematics are shown above timecourses. Box-and-whisker plots in **a**, **b**, and **d** show median, quartiles, min, and max. Data in **a**, **d**, and **e** analyzed using Mann-Whitney U-test. Maximum FRESCA and FRESCA2 response data in **a** and **b** reproduced from Fig. 1c.

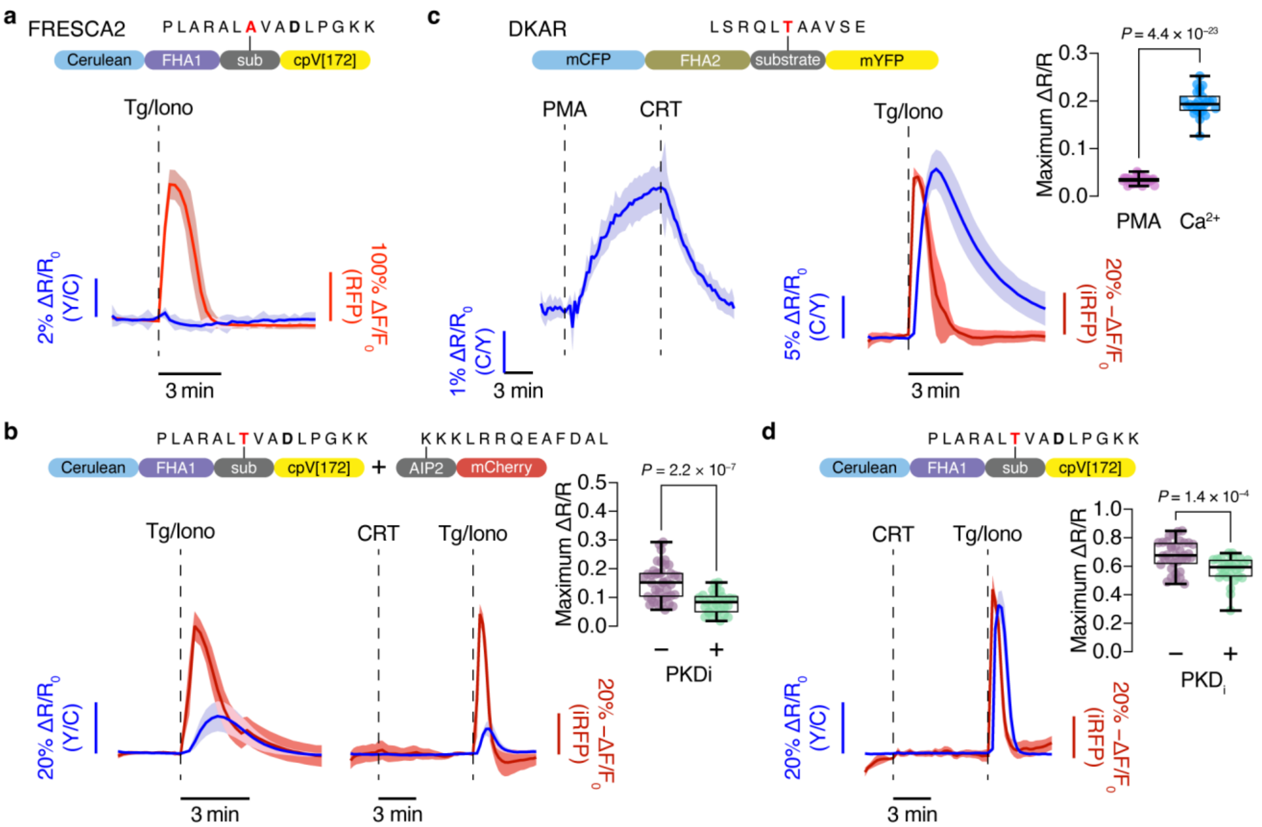


**Supplementary Figure 2 | FRESCA2 specificity in HeLa cells. a**, Representative average timecourse of FRESCA2 T/A ratio (blue) and RCaMP1d intensity (red) responses in Tg/Iono-stimulated HeLa cells. n=10 cells, representative of 3 independent experiments. **b**, PKD inhibition suppresses residual, CaMKII-inhibited FRESCA2 response. Representative average timecourse of emission ratio and fluorescence intensity responses from HeLa cells expressing FRESCA2 (blue curve) and NIR-GECO2G (dark red curve) plus CaMKip-mCherry and stimulated with Tg/Iono without (left, n=15 cells) or with (right, n=9 cells) CRT pretreatment. Data representative of 3 independent experiments each. Inset, maximum calcium-stimulated FRESCA2 ratio changes in HeLa cells co-expressing CaMKip-mCherry without (−; n=42 cells) or with (+; n=29 cells) CRT pretreatment. Data from 3 independent experiments each. **c**, Representative average timecourses of emission ratio and fluorescence intensity responses in HeLa cells expressing DKAR alone and stimulated with PMA and CRT (left, n=9 cells) or

DKAR plus NIR-GECO2G and stimulated with Tg/Iono (right, n=8 cells). Inset, maximum PMA (n=33 cells)- or calcium (n=26 cells)-stimulated DKAR emission ratio change. Data from 3 independent experiments each. **d**, Representative average timecourse of FRESCA2 ratio (blue) and NIR-GECO2G intensity (dark red) in HeLa cells treated with CRT followed by Tg/Iono. n=13 cells, representative of 3 independent experiments. Inset, maximum calcium-stimulated FRESCA2 emission ratio change without (−; n=36 cells) or with (+; n=28 cells) CRT pretreatment. Data from 3 independent experiments each. Solid lines in timecourse indicate the mean; shaded areas, s.d. Sensor schematics are shown above timecourses. Box-and-whisker plots inb, **c**, and **d** show median, quartiles, min, and max. Unpaired, two-tailed Student's t test without (**b**, **d**) or with (**d**) Welch's correction.

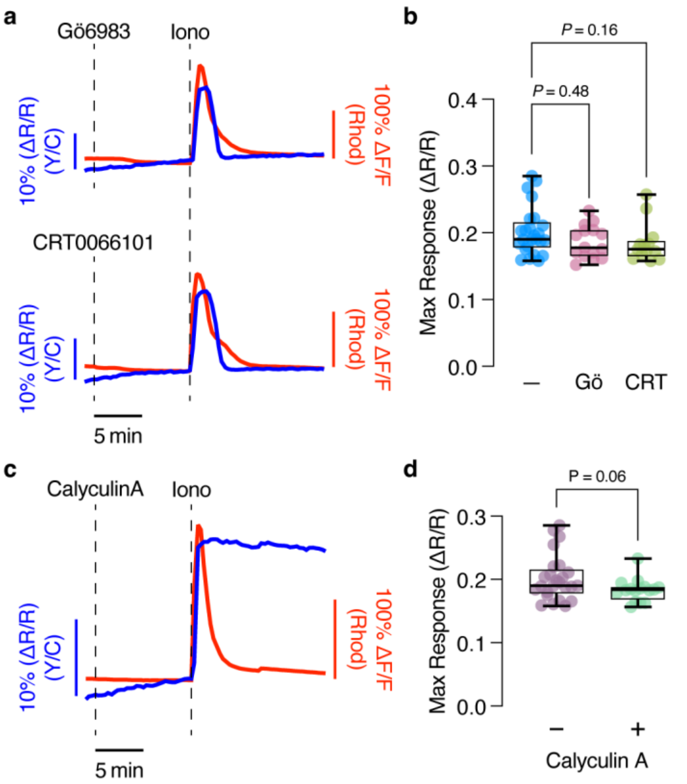


**Supplementary Figure 3 | Characterizing FRESCA2 in oocytes. a**, Representative timecourses showing the FRESCA2 ratio (blue) and Rhod-2 intensity (red) responses in oocytes stimulated with 500 nM Iono following pretreatment with either 1 μM Gö6983 (PKC inhibitor, top) or 5 μM CRT0066101 (PKD inhibitor, bottom). **b**, Maximum Iono-stimulated FRESCA2 emission ratio change (ΔR/R) in oocytes with or without PKC or PKD inhibition. n=25 (−), 15 (Gö), and 13 (CRT) eggs from 3 independent experiments. **c**, Phosphatase inhibition blocks reversal of the FRESCA2 response. Representative timecourse showing the FRESCA2 ratio (blue) and Rhod-2 intensity (red) in oocytes stimulated 500 nM Iono following pretreatment with 50 nM calyculin A. **d**, Maximum Iono-stimulated FRESCA2 emission ratio change (ΔR/R) in oocytes with or without calyculin A treatment. n=25 (−) and 16 (+) eggs from 4 and 3 independent experiments, respectively. Kruskal-Wallis test (**b**) or unpaired, two-tailed Student's t-test (**d**).

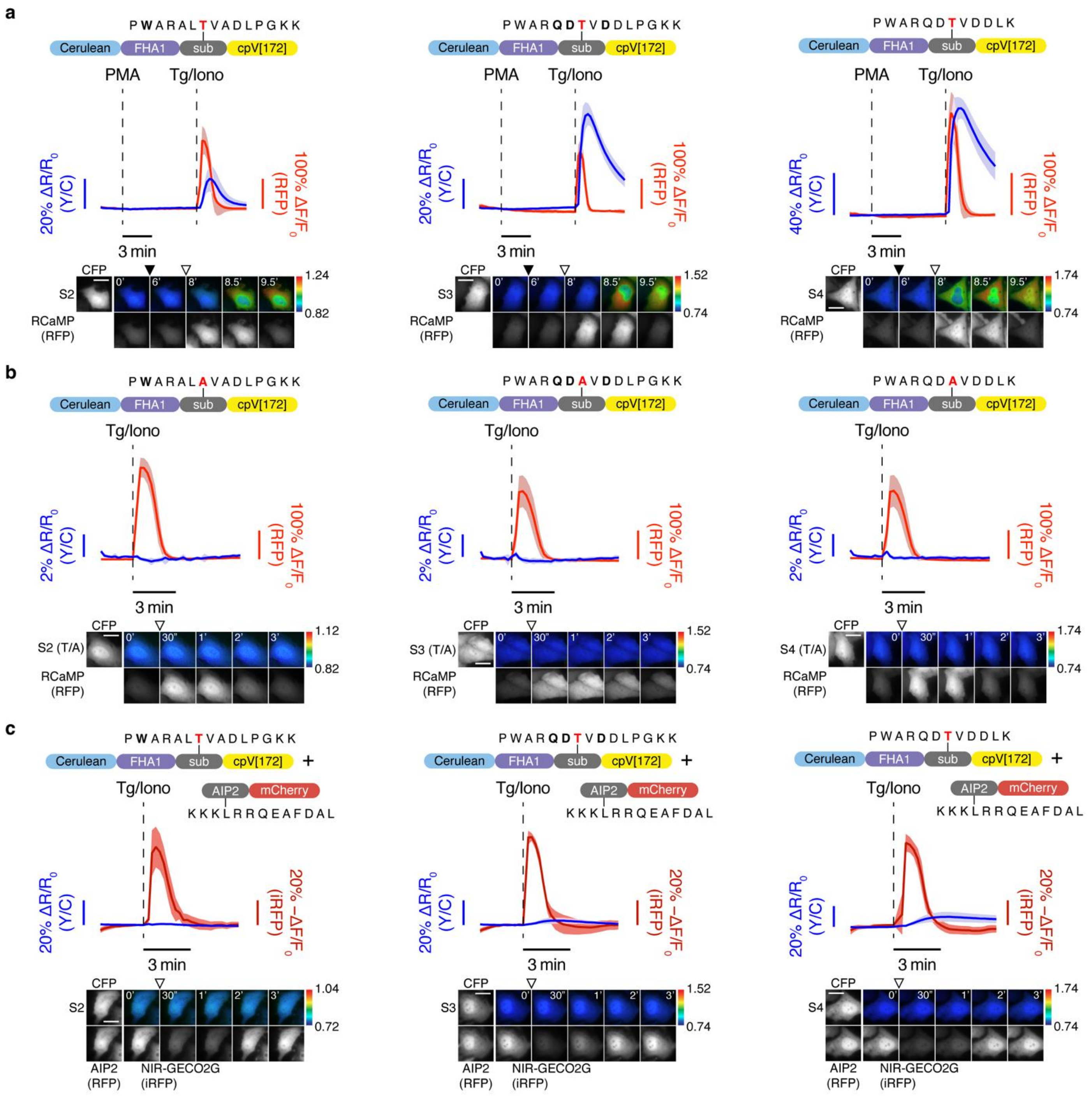


**Supplementary Figure 4 | Assessing CaMKII biosensor specificity. a**, Representative average timecourses showing ratio and intensity changes from (**a**) WT sensor variant (blue) S2 (top, n=9 cells), S3 (middle, n=10 cells), or S4 (bottom, n=9 cells) (**b**) T/A control sensor variant (blue curves) S2 (top, n=17 cells), S3 (middle, n=13 cells), or S4 (bottom, n=18 cells) and RCaMP1d (red), or (**c**) WT sensor variant (blue) S2 (top, n=11 cells), S3 (middle, n=8 cells), or S4 (bottom, n=9 cells) co-expressed with CaMKip-mCherry and NIR-GECO2G (dark red) in HeLa cells stimulated with Tg/Iono with (**a**) or without (**b**,**c**) PMA addition. All timecourses are representative of 3 independent experiments each. Solid lines indicate the mean; shaded areas, s.d. Sensor schematics are shown above timecourses. Images below timecourses show CaMKII sensor expression (CFP) and ratio (pseudocolor), along with RCaMP (RFP) intensity, before and after addition of PMA (closed arrowhead) and Tg/Iono (open arrowhead) as indicated. Scale bars, 20 μm.

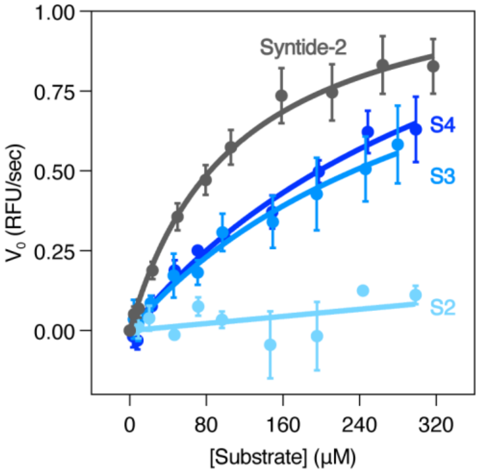


| Peptide | $V_{max}$ (RFU/s) | $K_M$ (µM) |
|---|---|---|
| Syntide-2 | 1.16 | 110.8 |
| S2 | — | — |
| S3 | 1.28 | 366.9 |
| S4 | 1.56 | 417.6 |

**Supplementary Figure 5 | *In vitro* phosphorylation of engineered CaMKII substrates.** Top, Michaelis-Menten plots or *in vitro* kinase reactions for different peptide substrates. Data points show mean ± s.e.m. from 3 independent experiments. Bottom, table summarizing fitted $V_{max}$ and $K_M$ values for each peptide substrate.

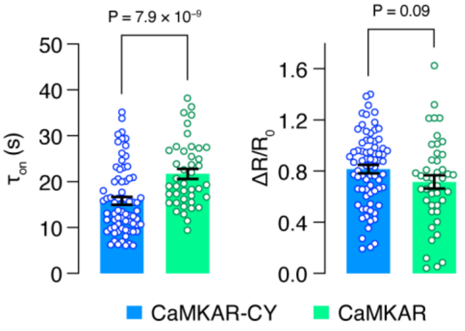


**Supplementary Figure 6 | CaMKAR-CY response shows rapid onset kinetics.** Quantification of $CaCl_2$-stimulated response onset kinetics ($\tau_{on}$, left) and maximum ratio change ($\Delta R/R_0$, right) in HEK-tsA201 cells expressing either CaMKAR-CY (blue, n=71 cells) or the previously reported excitation-ratiometric CaMKII sensor CaMKAR (green, n=42 cells) and stimulated as in Fig. 1a. Data from 3 independent experiments. Unpaired, two-tailed Student's t-tests.

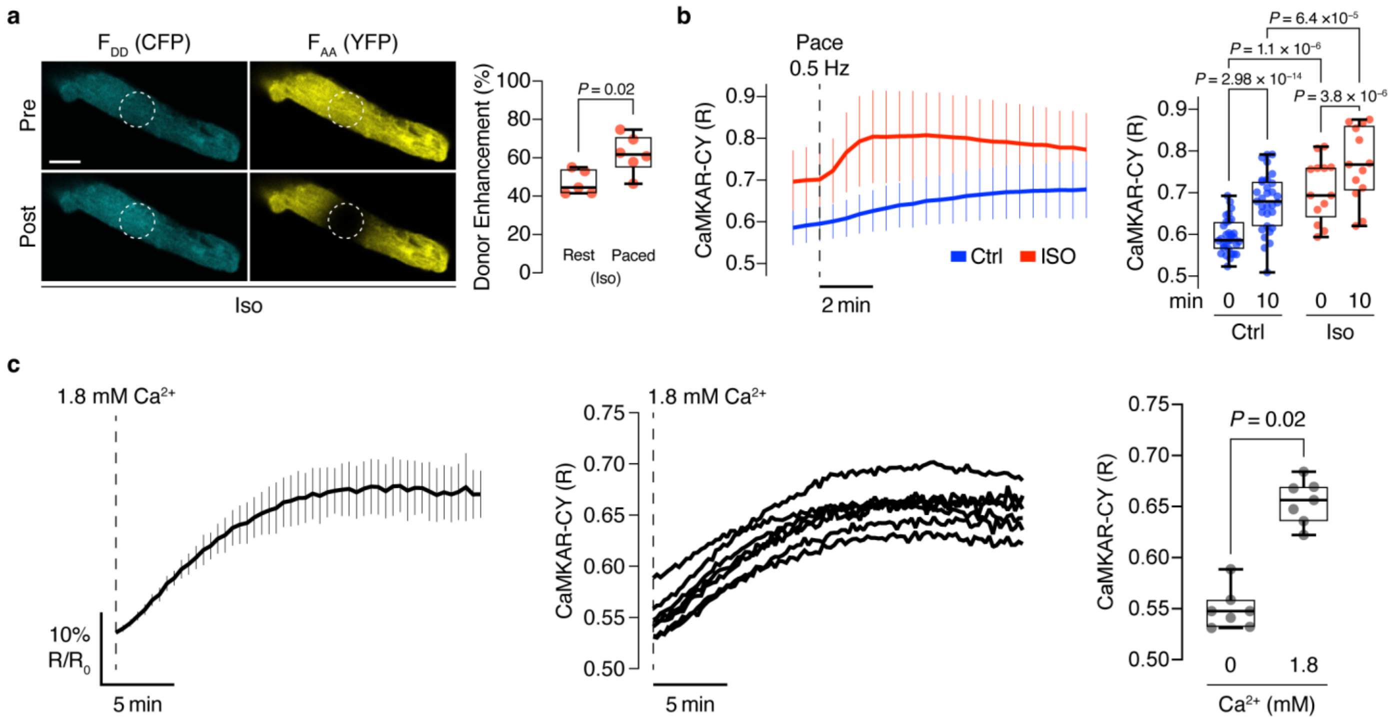


**Supplementary Figure 7 | CaMKAR-CY FRET verification and basal CaMKII activity in cardiac myocytes. a**, Verification of CaMKAR-CY FRET response in rabbit ventricular myocytes by donor enhancement upon acceptor photobleach. Left, representative images of an isoproterenol (Iso)-treated myocyte pre and post acceptor photobleach (>95% YFP bleach) resulting in enhanced donor (CFP) fluorescence. Scale bar, 20 μm. Right, quantification of donor enhancement (%) in Iso-treated (100 nM, incubation >3 min) myocytes at rest ($n_{rabbits}$=1; $n_{myocytes}$=5) vs. after 2 min of 1 Hz field-stimulation pacing ($n_{rabbits}$=1; $n_{myocytes}$=6). Mann-Whitney U test. **b**, Left, timecourses (mean ± s.d.) of averaged emission ratios ($R = F_{DA}/F_{DD}$) under Ctrl and Iso treatment conditions in response to 0.5 Hz field stimulation. Right, quantification of emission ratios (R) at 0 and 10 min after 0.5 Hz pacing onset under Ctrl and Iso treatment. Ctrl: $n_{rabbits}$=3, $n_{myocytes}$=31; Iso: $n_{rabbits}$=2, $n_{myocytes}$=13. **c**, Left, average timecourse showing normalized CaMKAR-CY ratio ($R/R_{0Ca}$) in resting ventricular myocytes pre-incubated (30 min) in $Ca^{2+}$-free solution upon raising $[Ca^{2+}]_o$ to 1.8 mM. Middle, individual nonnormalized R traces in response to $[Ca^{2+}]_o$ increase to 1.8 mM. The median R value at 0 mM $[Ca^{2+}]_o$, $R_{0Ca}$=0.548. Right, quantification of single-cell emission ratios (R). Solid lines in **b** and **c** indicate the mean; vertical lines, s.d. Wilcoxon matched-pairs signed rank test. Box-and-whisker plots in **b** and **c** show median, quartiles, min, and max. $n_{rabbits}$=1; $n_{myocytes}$=7.

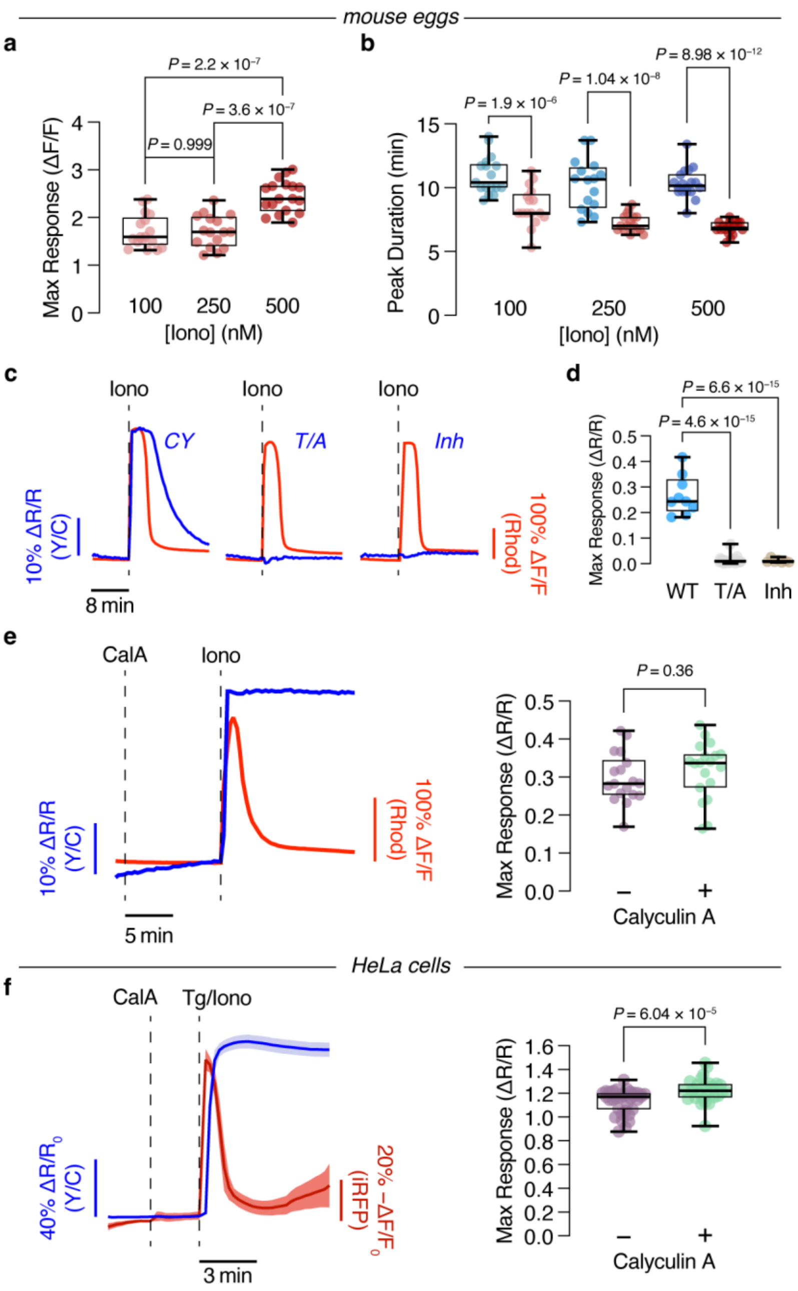


**Supplementary Figure 8 | Assessing CaMKAR-CY response in mouse oocytes. a**, Maximum Rhod-2 intensity change (ΔF/F) in oocytes stimulated with 100 nM, 250 nM, or 500 nM Iono. **b**, Duration of the Rhod and corresponding CaMKAR-CY response peaks at each Iono concentration. n=17 (100 nM), 16 (250 nM), and 18 (500 nM) eggs from 3 independent experiments each (both **a** and **b**). **c**, Representative traces of Y/C emission ratio (blue) and Rhod-

2 fluorescence intensity (red) in eggs expressing CaMKAR-CY (left), CaMKAR-CY T/A (middle), or CaMKAR-CY plus CaMKIIN1 (right) and stimulated with 500 nM Iono. Representative of n=9 (CY), 25 (T/A), and 7 (Inh) eggs from 3, 2, and 1 experiments, respectively. **d**, Quantification of maximum Iono-stimulated ratio change (ΔR/R) from eggs expressing CaMKAR-CY (WT), CaMKAR-CY T/A (T/A), or CaMKAR-CY plus CaMKIIN1 (Inh). n=9 (WT), 25 (T/A), and 7 (Inh) eggs from 3, 2, and 1 experiments, respectively. **e** and **f**, Phosphatase inhibition blocks CaMKAR-CY reversal. (**e**) Left: Representative timecourse showing CaMKAR-CY emission ratio (blue curve) and Rhod-2 fluorescence intensity (red curve) change in oocytes treated with 50 nM CalA prior to Iono stimulation. Representative of n=19 eggs. Right: Maximum Iono-stimulated ratio change (ΔR/R), with (n=18 eggs) or without (n=19 eggs) CalA pretreatment, from 4 independent experiments. (**f**) Left: Representative averaged timecourse showing CaMKAR-CY emission ratio (blue curve) and NIR-GECO2G fluorescence intensity (red curve) change in HeLa cells treated with 25 nM CalA prior to Tg/Iono stimulation (n=8 cells). Representative of 4 independent experiments. Right, summary of maximum CaMKAR-CY responses without (−; n=35 cells) or with (+; n=28 cells) CalA addition prior to Tg/Iono stimulation. Data are from 4 independent experiments.

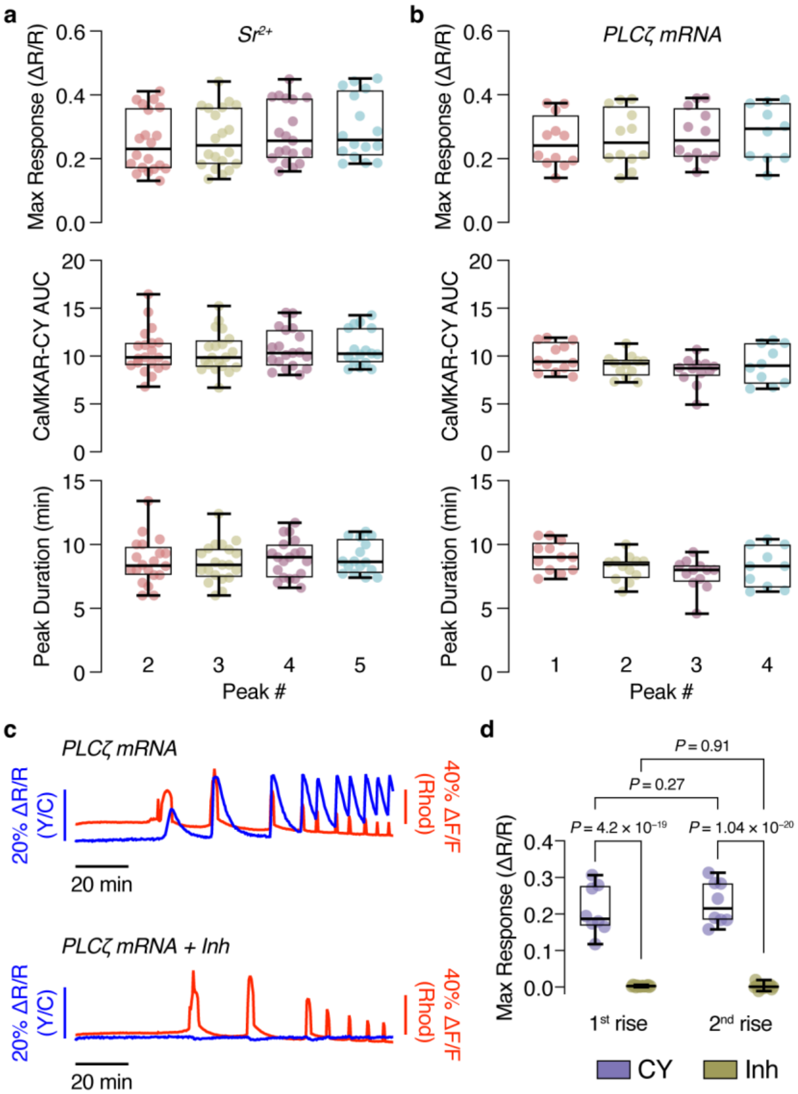


**Supplementary Figure 9 | Assessing CaMKAR-CY response in mouse oocytes. a** and **b**, Quantification of maximum CaMKAR-CY ratio change (ΔR/R), area under the curve (AUC), and peak duration for several consecutive transients during (**a**) $Sr^{2+}$- $Ca^{2+}$ oscillations or (**b**) PLCζ-induced $Ca^{2+}$ oscillations in mouse eggs. n=20 (**a**) and 12 (**b**) eggs from 4 independent experiments. **c**, Representative traces of CaMKAR-CY emission ratio (blue) and Rhod-2 fluorescence intensity (red) from mouse eggs injected with PLCζ mRNA without (upper) or with (lower) CaMKIIN1 mRNA (Inh). Representative of n=8 (CY) and 18 (Inh) eggs from 2 experiments each. **d**, Quantification of maximum PLCζ-induced CaMKAR-CY ratio change

(ΔR/R) without (CY) and with (Inh) CaMKIIN1 expression. n=8 (CY) and 18 (Inh) eggs from 2 experiments each.

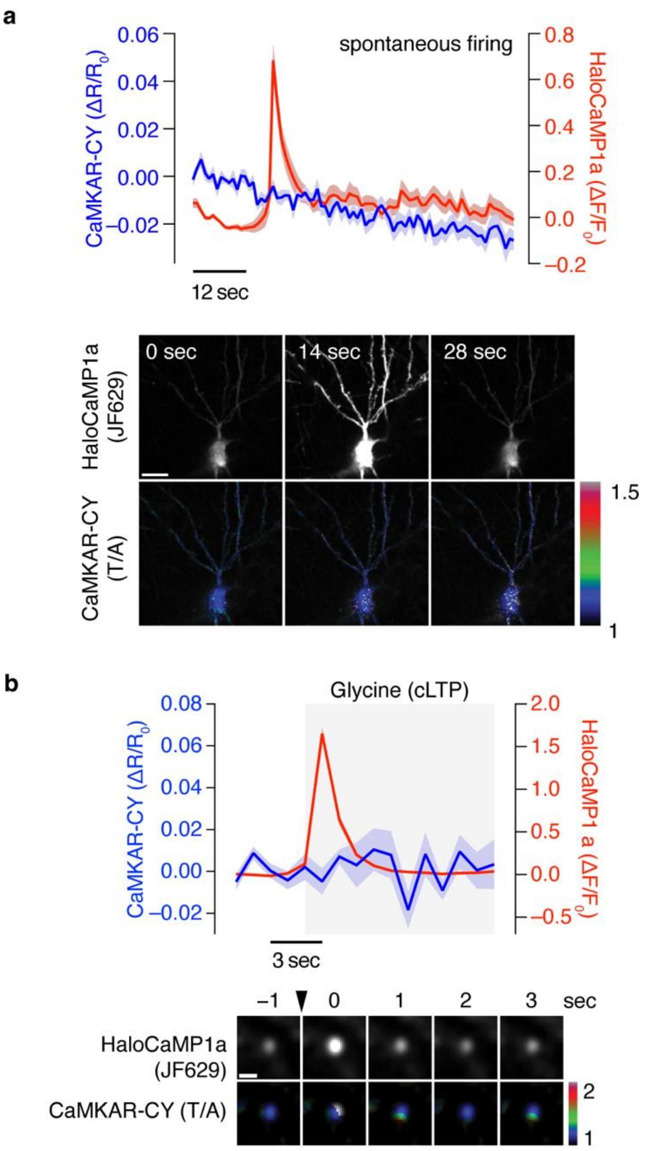


**Supplementary Figure 10 | CaMII activity imaging in hippocampal neurons using CaMKAR-CY. a**, Top, average responses from HaloCaMP1a (red trace) and CaMKAR-CY (T/A) (blue trace) aligned to $Ca^{2+}$ peaks from spontaneous firing events. n=37 cells from three experiments. Bottom, representative images of a spontaneous firing event (HaloCaMP, upper) and corresponding CaMKAR-CY (T/A) ratio images (lower). **b**, Top, average responses from HaloCaMP1a (red trace) and CaMKAR-CY (T/A) (blue trace) aligned to $Ca^{2+}$ peaks following cLTP induction. n=19 cells from three experiments. Bottom, representative images of an identified dendritic spine showing a $Ca^{2+}$ transient event (HaloCaMP, upper) following cLTP induction (arrowhead) and corresponding CaMKAR-CY (T/A) ratio images (lower). Solid lines in timecourses indicate means; shaded areas, s.e.m. Scale bars, 20 μm (**a**) and 2 μm (**b**).

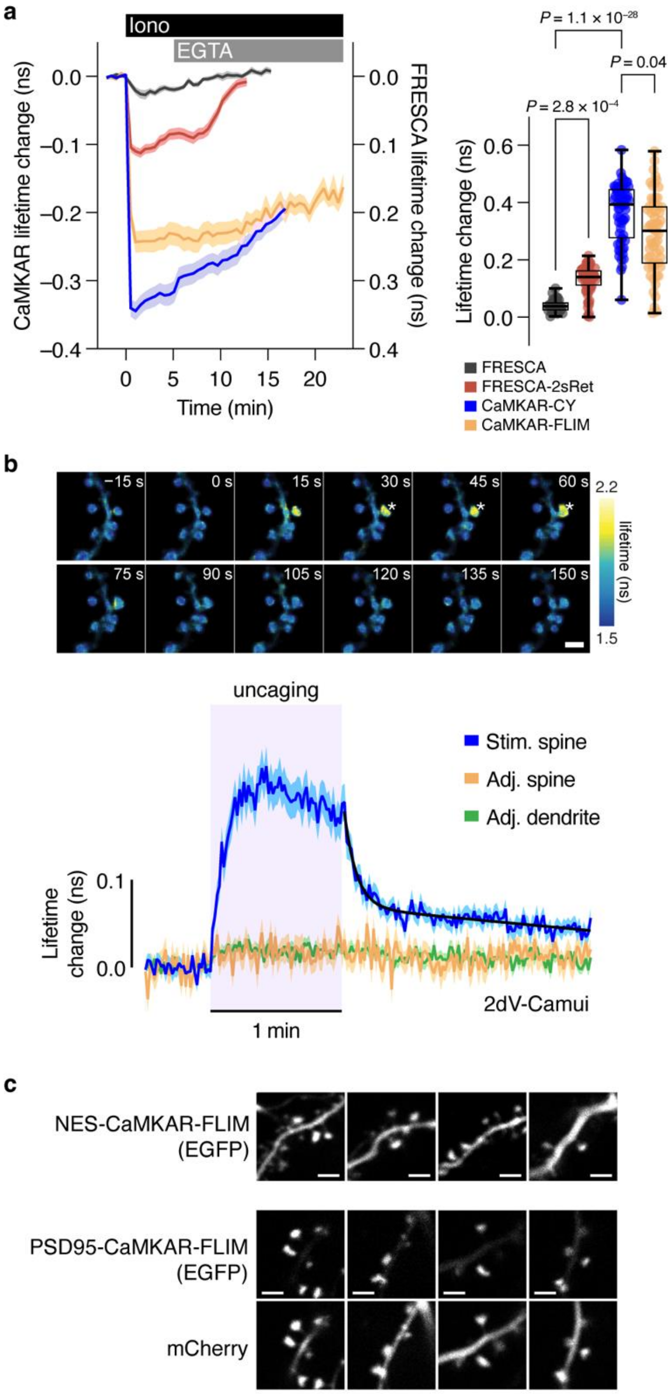


**Supplementary Figure 11 | CaMKII FLIM imaging in hippocampal CA1 synapses. a**, Left, averaged timecourses of fluorescence lifetime change of HEK cells transfected with indicated sensors plus CaMKIIα-IRES2-CaM (CaMKAR-CY, n=63; CaMKAR-FLIM, n=60; FRESCA, n=42; FRESCA-2sRet, n=67) upon treatment with ionomycin (12 μM) followed by EGTA (10 mM). Right, maximum fluorescence lifetime changes, plotted as absolute values for comparison. Box-and-whisker plots show median, quartiles, min, and max. Kruskal-Wallis test followed by Dunn's multiple-comparisons test. **b**, (top) Representative fluorescence lifetime images of dendritic spines during sLTP induction via glutamate uncaging in CA1 pyramidal neurons expressing 2dV-Camui in organotypic slice culture. Asterisks indicate the location of glutamate uncaging. Scale bar, 2 μm. (bottom) Average timecourses of fluorescence lifetime change of 2dV-Camui in the stimulated (Stim) spine, adjacent (Adj) spine (within ~7 μm of Stim spine), and adjacent dendrite (within ~4 μm of Stim spine). Black curve in shows double-exponential fit with time constants of 5.3 ± 0.6 s (62% fraction) and 224 ± 28 s (38% fraction). n (spines/neurons)=66/10. Solid lines and shaded areas in timecourses indicate mean and s.e.m., respectively. **c**, Representative EGFP fluorescence images of dendritic spines in CA1 pyramidal neurons expressing (top) NES-CaMKAR-FLIM or (bottom) PSD95-CaMKAR-FLIM highlighting sensor localization. Images of mCherry coexpressed as a morphology marker are also shown for PSD95-CaMKAR-FLIM-expressing neurons. Scale bars, 2 μm.

```
                     10         20         30         40         50         60         70
FRESCA2 (S1)  MVSKGEELFT GVVPILVELD GDVNGHRFSV SGEGEGDATY GKLTLKFICT TGKLPVPWPT LVTTLTWGVQ 70
          S2  MVSKGEELFT GVVPILVELD GDVNGHRFSV SGEGEGDATY GKLTLKFICT TGKLPVPWPT LVTTLTWGVQ 70
          S3  MVSKGEELFT GVVPILVELD GDVNGHRFSV SGEGEGDATY GKLTLKFICT TGKLPVPWPT LVTTLTWGVQ 70
 CaMKAR-CY (S4)  MVSKGEELFT GVVPILVELD GDVNGHRFSV SGEGEGDATY GKLTLKFICT TGKLPVPWPT LVTTLTWGVQ 70
CaMKAR-FLIM (S4)  MVSKGEELFT GVVPILVELD GDVNGHKFSV SGEGEGDATY GKLTLKFICT TGKLPVPWPT LVTTLTYGVQ 70

                     80         90        100        110        120        130        140
FRESCA2 (S1)  CFARYPDHMK QHDFFKSAMP EGYVQERTIF FKDDGNYKTR AEVKFEGDTL VNRIELKGID FKEDGNILGH 140
          S2  CFARYPDHMK QHDFFKSAMP EGYVQERTIF FKDDGNYKTR AEVKFEGDTL VNRIELKGID FKEDGNILGH 140
          S3  CFARYPDHMK QHDFFKSAMP EGYVQERTIF FKDDGNYKTR AEVKFEGDTL VNRIELKGID FKEDGNILGH 140
 CaMKAR-CY (S4)  CFARYPDHMK QHDFFKSAMP EGYVQERTIF FKDDGNYKTR AEVKFEGDTL VNRIELKGID FKEDGNILGH 140
CaMKAR-FLIM (S4)  CFSRYPDHMK QHDFFKSAMP EGYVQERTIF FKDDGNYKTR AEVKFEGDTL VNRIELKGID FKEDGNILGH 140

                    150        160        170        180        190        200        210
FRESCA2 (S1)  KLEYNAISDN VYITADKQKN GIKAHFKIRH NIEDGSVQLA DHYQQNTPIG DGPVLLPDNH YLSTQSALSK 210
          S2  KLEYNAISDN VYITADKQKN GIKAHFKIRH NIEDGSVQLA DHYQQNTPIG DGPVLLPDNH YLSTQSALSK 210
          S3  KLEYNAISDN VYITADKQKN GIKAHFKIRH NIEDGSVQLA DHYQQNTPIG DGPVLLPDNH YLSTQSALSK 210
 CaMKAR-CY (S4)  KLEYNAISDN VYITADKQKN GIKAHFKIRH NIEDGSVQLA DHYQQNTPIG DGPVLLPDNH YLSTQSALSK 210
CaMKAR-FLIM (S4)  KLEYNYNSHN VYIMADKQKN GIKANFKIRH NIEDGSVQLA DHYQQNTPIG DGPVLLPDNH YLSTQSALSK 210

                    220        230        240        250        260        270        280
FRESCA2 (S1)  DPNEKRDHMV LLEFVTAAGI TLGMDELYKR MHKFSQEQIG ENIVCRVICT TGQIPIRDLS ADISQVLKEK 280
          S2  DPNEKRDHMV LLEFVTAAGI TLGMDELYKR MHKFSQEQIG ENIVCRVICT TGQIPIRDLS ADISQVLKEK 280
          S3  DPNEKRDHMV LLEFVTAAGI TLGMDELYKR MHKFSQEQIG ENIVCRVICT TGQIPIRDLS ADISQVLKEK 280
 CaMKAR-CY (S4)  DPNEKRDHMV LLEFVTAAGI TLGMDELYKR MHKFSQEQIG ENIVCRVICT TGQIPIRDLS ADISQVLKEK 280
CaMKAR-FLIM (S4)  DPNEKRDHMV LLEFVTAAGI TLGMDELYKR MHKFSQEQIG ENIVCRVICT TGQIPIRDLS ADISQVLKEK 280

                    290        300        310        320        330        340        350
FRESCA2 (S1)  RSIKKVWTFG RNPACDYHLG NISRLSNKHF QILLGEDGNL LLNDISTNGT WLNGQKVEKN SNQLLSQGDE 350
          S2  RSIKKVWTFG RNPACDYHLG NISRLSNKHF QILLGEDGNL LLNDISTNGT WLNGQKVEKN SNQLLSQGDE 350
          S3  RSIKKVWTFG RNPACDYHLG NISRLSNKHF QILLGEDGNL LLNDISTNGT WLNGQKVEKN SNQLLSQGDE 350
 CaMKAR-CY (S4)  RSIKKVWTFG RNPACDYHLG NISRLSNKHF QILLGEDGNL LLNDISTNGT WLNGQKVEKN SNQLLSQGDE 350
CaMKAR-FLIM (S4)  RSIKKVWTFG RNPACDYHLG NISRLSNKHF QILLGEDGNL LLNDISTNGT WLNGQKVEKN SNQLLSQGDE 350

                    360        370        380        390        400        410        420
FRESCA2 (S1)  ITVGVGVESD ILSLVIFIND KFKQCLEQNK VDRSAGKPGS GEGSTKGPLA RALTVADLPG KKGGTGGSEL 420
          S2  ITVGVGVESD ILSLVIFIND KFKQCLEQNK VDRSAGKPGS GEGSTKGPWA RALTVADLPG KKGGTGGSEL 420
          S3  ITVGVGVESD ILSLVIFIND KFKQCLEQNK VDRSAGKPGS GEGSTKGPWA RQDTVDDLPG KKGGTGGSEL 420
 CaMKAR-CY (S4)  ITVGVGVESD ILSLVIFIND KFKQCLEQNK VDRSAGKPGS GEGSTKGPWA RQDTVDDL-- -KGGTGGSEL 417
CaMKAR-FLIM (S4)  ITVGVGVESD ILSLVIFIND KFKQCLEQNK VDRSAGKPGS GEGSTKGPWA RQDTVDDL-- -KGGTGGSEL 417

                    430        440        450        460        470        480        490
FRESCA2 (S1)  MGGVQLADHY QQNTPIGDGP VLLPDNHYL SYQSKLSKDPN EKRDHMVLLE FVTAAGITLG MDELYKGGTG 490
          S2  MGGVQLADHY QQNTPIGDGP VLLPDNHYL SYQSKLSKDPN EKRDHMVLLE FVTAAGITLG MDELYKGGTG 490
          S3  MGGVQLADHY QQNTPIGDGP VLLPDNHYL SYQSKLSKDPN EKRDHMVLLE FVTAAGITLG MDELYKGGTG 490
 CaMKAR-CY (S4)  MGGVQLADHY QQNTPIGDGP VLLPDNHYL SYQSKLSKDPN EKRDHMVLLE FVTAAGITLG MDELYKGGTG 487
CaMKAR-FLIM (S4)  MGSVQLADHY QQNTPIGDGP VLLPDNHYLS YQSALSKDPN EKRDHMVLLE RVTAAGITLG MDELYKGGTG 487

                    500        510        520        530        540        550        560
FRESCA2 (S1)  GSMVSKGEEL FTGVVPILVE LDGDVNGHKF SVSGEGEGDA TYGKLTLKLI CTTGKLPVPW PTLVTTLGYG 560
          S2  GSMVSKGEEL FTGVVPILVE LDGDVNGHKF SVSGEGEGDA TYGKLTLKLI CTTGKLPVPW PTLVTTLGYG 560
          S3  GSMVSKGEEL FTGVVPILVE LDGDVNGHKF SVSGEGEGDA TYGKLTLKLI CTTGKLPVPW PTLVTTLGYG 560
 CaMKAR-CY (S4)  GSMVSKGEEL FTGVVPILVE LDGDVNGHKF SVSGEGEGDA TYGKLTLKLI CTTGKLPVPW PTLVTTLGYG 557
CaMKAR-FLIM (S4)  GSMVSKGEEL FTGVVPILVE LDGDVNGHKF SVSGEGEGDA TYGKLTLKLI CTTGKLPVPW PTLVTTFGYG 557

                    570        580        590        600        610        620        630
FRESCA2 (S1)  LQCFARYPDH MKQHDFFKSA MPEGYVQERT IFFKDDGNYK TRAEVKFEGD TLVNRIELKG IDFKEDGNIL 630
          S2  LQCFARYPDH MKQHDFFKSA MPEGYVQERT IFFKDDGNYK TRAEVKFEGD TLVNRIELKG IDFKEDGNIL 630
          S3  LQCFARYPDH MKQHDFFKSA MPEGYVQERT IFFKDDGNYK TRAEVKFEGD TLVNRIELKG IDFKEDGNIL 630
 CaMKAR-CY (S4)  LQCFARYPDH MKQHDFFKSA MPEGYVQERT IFFKDDGNYK TRAEVKFEGD TLVNRIELKG IDFKEDGNIL 627
CaMKAR-FLIM (S4)  LMCFARYPDH MKQHDFFKSA MPEGYVQERT IFFKDDGNYK TRAEVKFEGD TLVNRIELKG IDFKEDGNIL 627

                    640        650        660
FRESCA2 (S1)  GHKLEYNYNS HNVYITADKQ KNGIKANFKI RHNIE 665
          S2  GHKLEYNYNS HNVYITADKQ KNGIKANFKI RHNIE 665
          S3  GHKLEYNYNS HNVYITADKQ KNGIKANFKI RHNIE 665
 CaMKAR-CY (S4)  GHKLEYNYNS HNVYITADKQ KNGIKANFKI RHNIE 662
CaMKAR-FLIM (S4)  GHKLEYNWNS VNVYIMADKQ KNGIKVNFKI RHNIE 662
```

**Supplementary Figure 12. Primary sequences of engineered CaMKII biosensors.** Color scheme corresponds to biosensor schematics shown in Figure 1b and Figure 6a. Substrate modifications in S2-S4 are indicated in red.

### Supplementary Tables

**Supplementary Table 1. CaMKII substrate scoring summary.** Summary of Log2 scores and score ranks for CAMK2A and PRKD1 phosphorylation of the indicated sequences, obtained from https://www.phosphosite.org/kinaseLibraryAction using default settings. For each sequence, only the scorable region ($P_{-5}$ to $P_{+4}$) is shown, with the phosphosite ($P_0$) denoted by *.

| | CAMK2A | | PRKD1 | |
|---|---|---|---|---|
| **Sequence** | **Log2 Score** | **Score Rank** | **Log2 Score** | **Score Rank** |
| S1 – LARALT*VADL | 1.97 | 38 | 3.30 | 22 |
| S2 – WARALT*VADL | 1.79 | 38 | 0.64 | 83 |
| S3/S4 – WARQDT*VDDL | 7.17 | 1 | 1.00 | 41 |
